# Microbial metabolic potential of hydrothermal vent chimneys along the Submarine Ring of Fire

**DOI:** 10.1101/2023.09.12.557424

**Authors:** Laura Murray, Heather Fullerton, Craig L. Moyer

**Affiliations:** Department of Biology, Western Washington University, Bellingham, WA, United States; Department of Biology, College of Charleston, Charleston, SC, United States

**Keywords:** hydrothermal vents, chimneys, community structure, metagenomics, metabolism

## Abstract

Hydrothermal vents host a diverse community of microorganisms that utilize chemical gradients from the venting fluid for their metabolisms. The venting fluid can solidify to form chimney structures that these microbes adhere to and colonize. These chimney structures are found throughout many different locations in the world’s oceans. In this study, comparative metagenomic analyses of microbial communities on five chimney structures from around the Pacific Ocean were elucidated focusing on the core taxa and genes that are characteristic for each of these hydrothermal vent chimneys, as well as highlighting differences among the taxa and genes found at each chimney due to parameters such as physical characteristics, chemistry, and activity of the vents. DNA from the chimneys was sequenced, assembled into contigs, annotated for gene function, and binned into metagenome-assembled genomes, or MAGs. Genes used for carbon, oxygen, sulfur, nitrogen, iron, and arsenic metabolism were found at varying abundances at each of the chimneys, largely from either Gammaproteobacteria or Campylobacteria. Many taxa had an overlap of these metabolic genes, indicating that functional redundancy is critical for life at these hydrothermal vents. A high relative abundance of oxygen metabolism genes coupled with low carbon fixation genes could be used as a unique identifier for inactive chimneys. Genes used for DNA repair, chemotaxis, and transposases were found to be at higher abundances at each of these hydrothermal chimneys allowing for enhanced adaptations to the ever-changing chemical and physical conditions encountered.

**IMPORTANCE:** The metabolic byproducts of microorganisms that form and reside in hydrothermal vent chimneys facilitate nutrient cycling in both the hydrothermal vent ecosystem and throughout the world’s oceans. Diverse communities of microbes utilize chemicals in the venting fluid to gain energy and biomass. Here, metagenomic and amplicon sequencing was used to identify metabolism genes to better understand the metabolic potential of chimneys. The combination of genes detected in this study sheds light on hydrothermal vent chimneys’ community structure and metabolic potential throughout the Pacific Ocean.

## INTRODUCTION

Deep-sea hydrothermal vents have been an ecosystem of interest since their discovery in 1977 (1). The mixing of the hot reduced fluids with the cold oxygenated seawater creates the chimney, solid structures in which organisms can adhere to (2). Not only do these chimneys provide a surface for microbial growth, but vent fluids support the growth of chemoautotrophic microbes (3). The metabolic byproducts of these microbes are disseminated throughout the ocean; contributing to global geochemical cycling and utilized as energy sources by macroorganisms throughout the world’s oceans (4, 5). By metabolizing vent fluid and mineral deposits of the hydrothermal vent chimneys, chemoautotrophic microbes enhance community richness and diversity by allowing for hydrothermal vents to become more diverse communities (6).

Hydrothermal vents are ecologically and economically important, due to the high levels of valuable metals like silver, copper, cobalt, and gold (7). These ecosystems are becoming popular sites for deep-sea mining due to the concentrations of metals. Deep-sea mining produces large particle plumes and landslides that can impact benthic filter feeders’ ability to access food and destroy the growth surfaces (8). This could have downstream impacts since these microbes could act as settlement cues for vent-endemic invertebrates (9). Without these microbial primary producers at these chimney sites, the rich ecosystem would become largely uninhabitable.

Chimneys are found worldwide and have a wide range of venting temperatures, chemical composition, and flow rates (10). Differences in abiotic factors, such as the presence of oxygen, pH, pressure, heat flux, and reduced chemicals dictate the biogeographical distribution of microbes. However, due to the necessity of microbes having optimal conditions needed for growth, vents with similar environmental characteristics have similar microbial communities (11).

Magic Mountain has over 50 active and inactive vents, including the inactive Ochre Chimney. Active and inactive chimney structures found in this vent field are composed largely of sulfide deposits (12). Active chimneys at Magic Mountain are dominated by Campylobacteria, (formerly known as Epsilonproteobacteria (13)). When examining inactive chimneys in other locations, sulfide-oxidizing Gammaproteobacteria tend to dominate the microbial communities as they can use metal-sulfide present in the chimney structures and mineral deposits for energy (14–16).

Axial Seamount is an active submarine volcano and a site of extensive hydrothermal venting, with the most recent eruption occurring in 2015 (17). The high volcanic activity of this caldera produces a large amount of hydrogen sulfide, ferrous oxide, and methane that are released during eruption events. An abundance of Sulfur-oxidizing bacteria and methanogenic archaea correlate to the high concentrations of hydrogen sulfide and methane (2). Axial Seamount plumes and microbial mats showed Aquificae, Gammaproteobacteria, Campylobacteria, and classes of methanogenic archaea dominated the microbial communities. The geochemistry of the venting sites is important in shaping community structure and genes present at each vent (18).

The Guaymas Basin is highly active with steep temperature gradients and is covered with a layer of a few hundred meters of organic-rich sediment. Some chimneys here are characterized not by direct venting but by internal hydrothermal fluid circulation due to their shape (19). These unique chimneys have pagoda-like structures that have fluid circulating internally, leading to large, temperature gradients. Metagenomic analysis of Guaymas Basin sediments showed an enrichment of genes for methane, hydrogen, and sulfide metabolisms as compared to background sediments (20). Whereas, metagenomic analysis of a chimney sample demonstrated that heterotrophic sulfate-reducing bacteria were found at higher abundances due to the high concentrations of hydrocarbons found in the Guaymas Basin (21).

At Urashima, dissolved sulfide and hydrogen concentrations in vent effluent are enhanced. Due to the high levels of sulfide, the microbial community is dominated by the sulfur oxidizer Campylobacteria (22). The Urashima Vent Field has shown to have a relatively high abundance of Zetaproteobacteria, a class of iron-oxidizing bacteria that forms dense mat structures made from iron oxides and polysaccharides which are used as a colonizing surface for other microbial taxa (23).

In this study, we seek to characterize the microbial community members and metabolic potential at each chimney found along the plate boundary in the Pacific Ocean, known as the “Submarine Ring of Fire” (Figure 1). This approach will allow for investigations into the similarities and differences in microbial community structure and identification of the common genes found at each chimney across these selected vent chimneys.

**Figure 1.**
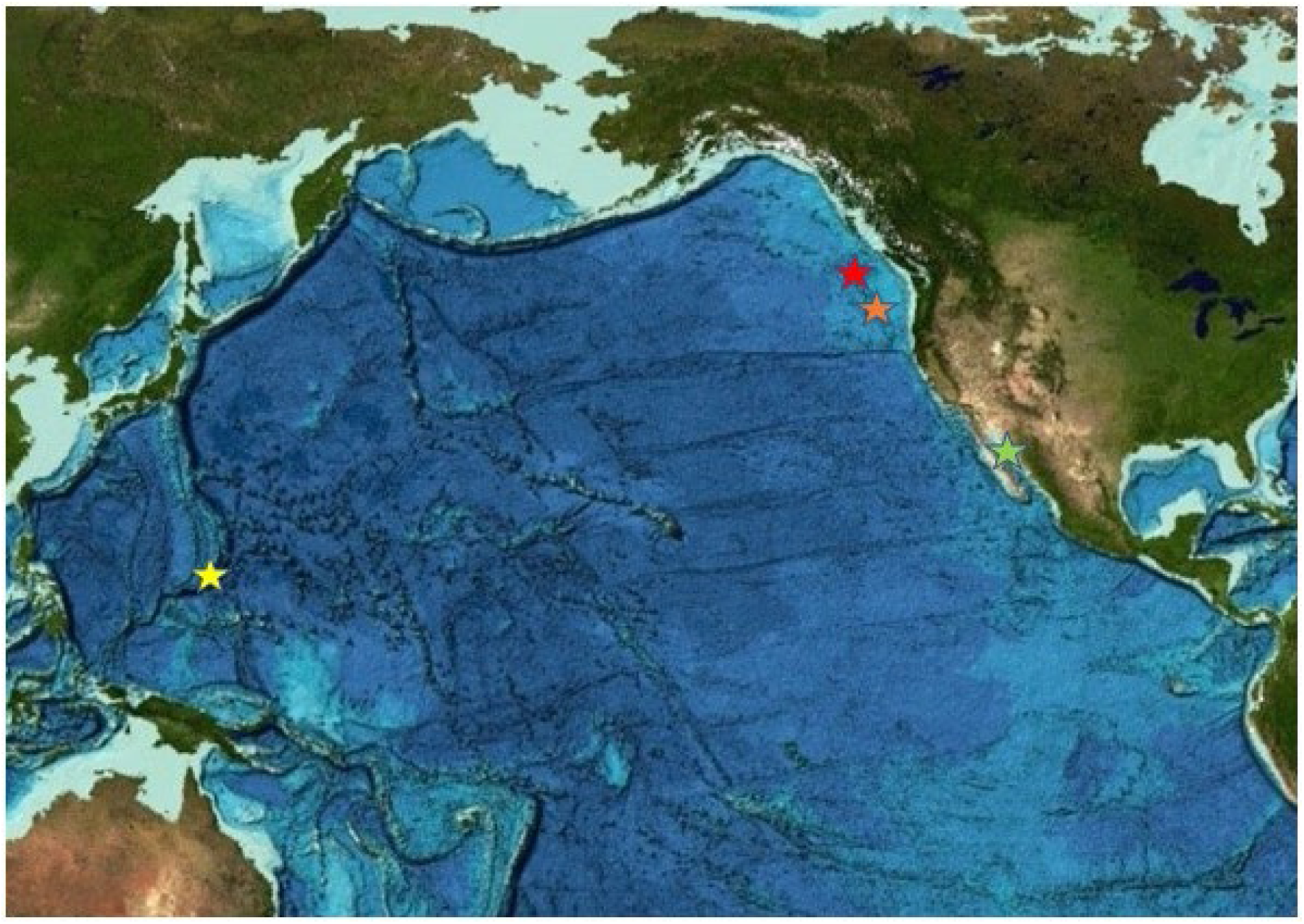
Map of the four sampling locations across the Pacific Ocean. The red star denotes the Magic Mountain, Explorer Ridge sampling location of the Ochre Chimney, the orange star denotes the Axial Volcano, Juan de Fuca Ridge sampling location of the Castle Chimney. The green star denotes the Guaymas Basin sampling location of the Pagoda Chimney. The yellow star denotes the Mariana back-arc Urashima sampling location of the Snap-Snap and Ultra-No-Chi-Chi Chimneys. (Image reproduced from the GEBCO world map 2019, www.gebco.net.)

**Figure 2.**
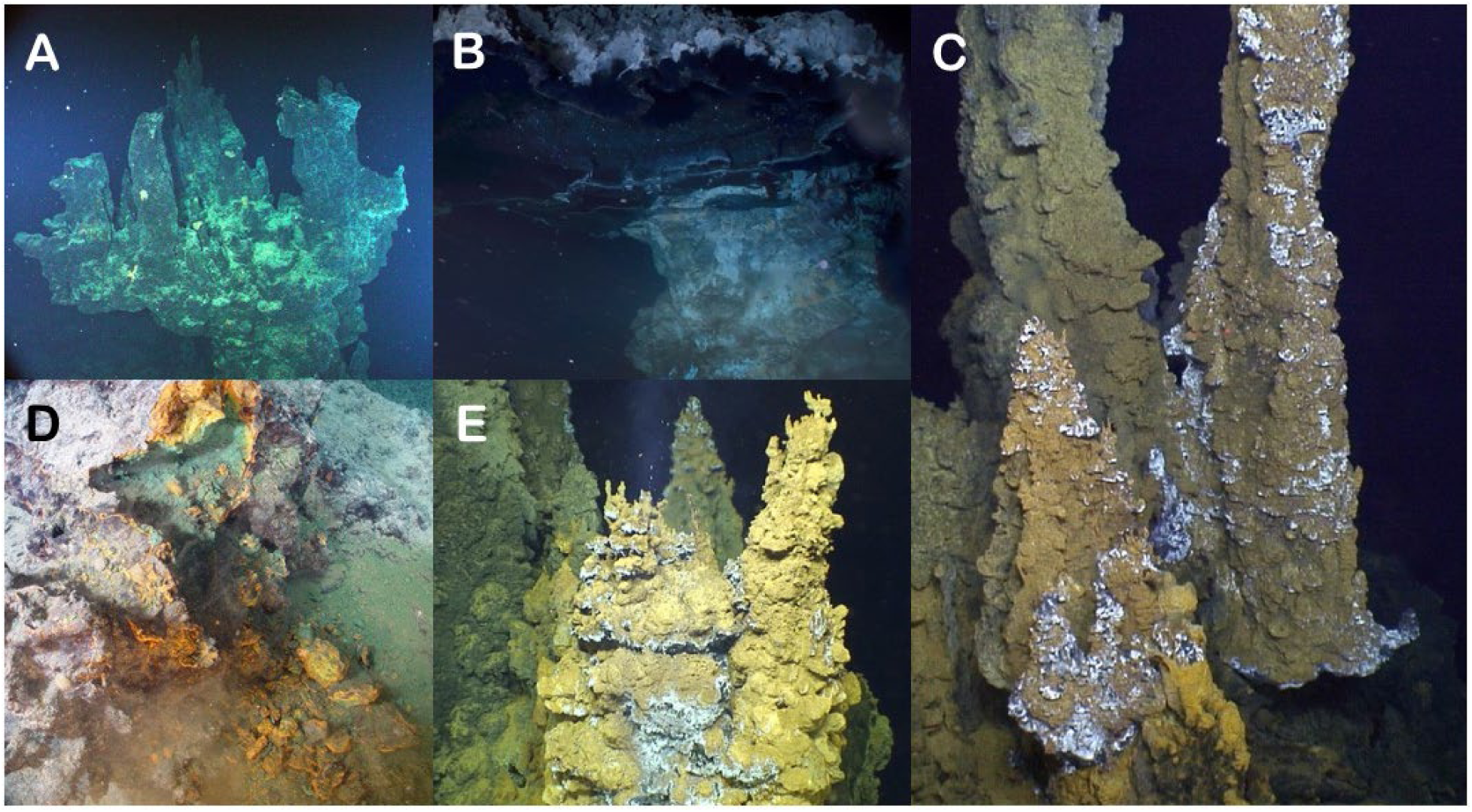
Photos of the five chimneys evaluated in this study. (A) Castle Chimney from Axial Seamount. (B) Pagoda Chimney from Guaymas Basin. (C) Ultra-No-Chi-Chi Chimney from the Urashima Vent Field (Laser dots are 10 cm apart). (D) Ochre Chimney from Magic Mountain. (E) Snap-Snap Chimney also from the Urashima Vent Field.

## RESULTS

### Assembly

After sequencing and filtration for quality, 207,450,568 reads were assembled into 15,601,568 contigs. The N50 was 385 base pairs which had an average coverage of 0.669 at Ochre, 0.815 at Castle, 0.515 at Pagoda, 0.388 at Snap, and 0.446 at Ultra. After composite assembly, a total of 17,978,241 ORFs were identified and of these 40.7% were able to be annotated. The composite assembly resulted in 43 MAGs with a completeness range of 10% to 52% and a contamination range of 0% to 4.76% (Supp. Table 1). Due to the low completeness and high contamination level of the MAGs, further analysis was done directly on the composite assembly.

### Alpha and Beta Diversity

All five chimneys were distinctly separated in the NMDS plot, indicating unique environmental parameters at each chimney (Figure 3A). Clustering of samples using Bray-Curtis distance matrices and NMDS plot of KEGG gene functions showed the inactive Ochre Chimney further separated from the other chimney samples and that the two Urashima chimneys cluster closest together (Figure 3A). When clustered by taxa, Pagoda Chimney clustered as an outgroup, while the two Urashima chimneys clustered most closely together, and the Castle and Ochre chimneys also clustered together (Figure 3B). The amplicon sequence analysis showed the two Urashima chimneys cluster most closely together, similar to the SSU rRNA genes from the composite assembly (Figure 3C). However, with the amplicon analysis, Ochre Chimney was the outgroup. Despite each chimney having a significant R^2^ value after PERMANOVA analysis, the p-value was 0.1 or greater, likely due to each chimney only having a single sample.

**Figure 3.**
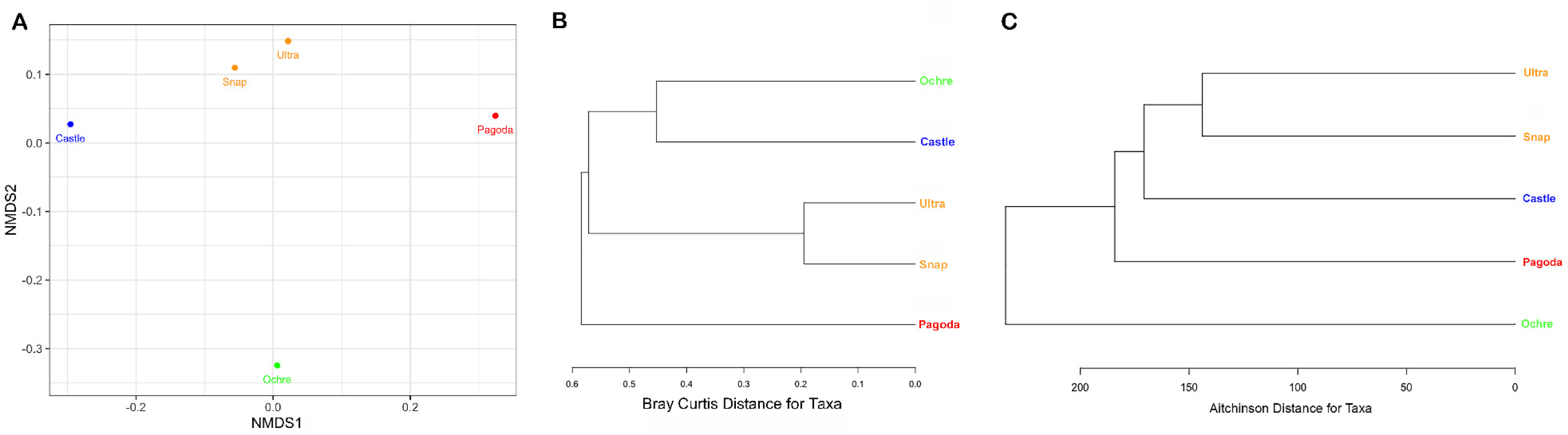
(A) NMDS plot using Bray-Curtis distance metric with the chimneys labeled with colored dots by location. The blue dot represents Castle Chimney at Axial Seamount, green represents Ochre Chimney at Magic Mountain, red represents Pagoda Chimney at Guaymas Basin Vent Field and orange represents the Snap-Snap and Ultra-No-Chi-Chi Chimneys from the Urashima Vent Field. (B) Cluster analysis of metagenomic-derived taxa similarities among chimneys using Bray Curtis distance. (C) Cluster analysis of amplicon sequencing-derived taxa similarities among chimneys using Aitchinson distance.

Each chimney had variations in the taxonomic identity of ORF, the identity of SSU rRNA genes in the composite assembly, and SSU rRNA amplicon identity (Figure 4). The composite metagenomes resulted in the identification of 23,040 SSU rRNA gene whereas amplicon analysis resulted in 5,380 ASVs. By ORF analysis, the Castle Chimney community was composed of 38% unclassified Bacteria and 38% Gammaproteobacteria, with 15% unclassified Proteobacteria, (newly named to Pseudomonadota (24)), with less than 10% Alphaproteobacteria, unclassified Bacteroidetes, and Deltaproteobacteria. These were identified as 60% unclassified Bacteria and 40% Gammaproteobacteria (Fig. 4B). By amplicon sequencing, Castle had 50% Gammaproteobacteria, 20% Thermodesulfovibriornia, 15% Bacteroidia, 10% Alphaproteobacteria and 5% Ignavibacteria (Figure 4C).

**Figure 4.**
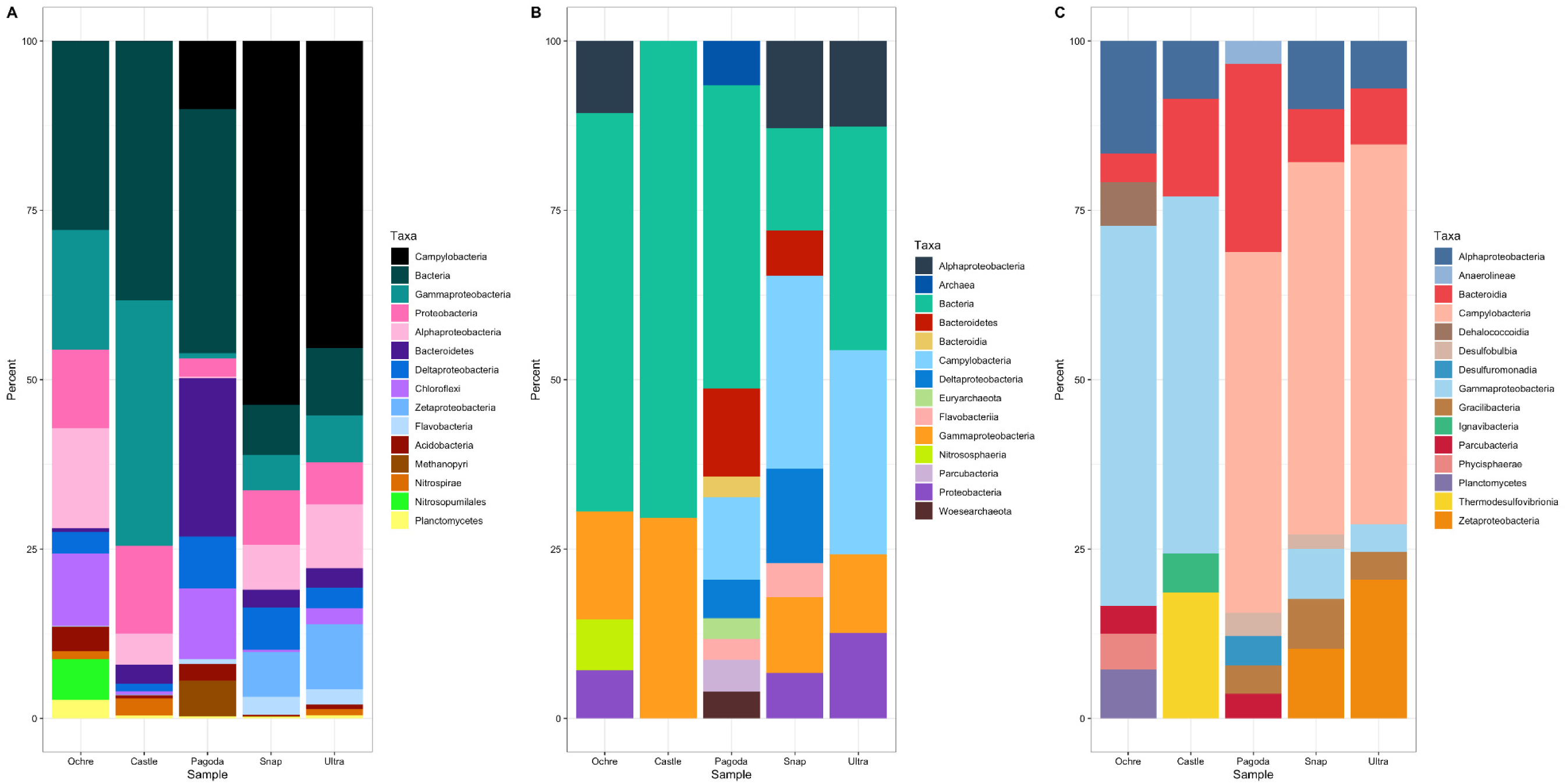
Stacked bar graph of the top 15 microbial taxa found in each chimney as a percentage of reads from the whole metagenome and from all taxa detected by amplicon sequencing. (A) Stacked bar graph of the ORFs from assembly, not including unclassified or unmapped reads. (B) Stacked bar graph of the SSU rRNA gene reads, only including the top 15 taxa, not including unclassified or unmapped metagenome reads. (C) Stacked bar graph based on amplicon sequencing of SSU rRNA genes using V3V4 primers.

Pagoda Chimney’s ORFs were comprised of 30% unclassified Bacteria and 21% unclassified Bacteroidetes, with 10% Campylobacteria, 10% unclassified Chloroflexi 9% Deltaproteobacteria, and 5% Methanopyri. By SSU rRNA genes of the composite assembly, Pagoda had 35% unclassified Bacteria, 10% Bacteroidetes, 10% Campylobacteria, and less than 10% of unclassified Archaea, Bacteroidia, Deltaporotebacteria, Parcubacteria, Euryarchaeota, Flavobacteriia, and Woesarchaeota (Figure 4B). Amplicon sequencing showed 25% Bacteroidia, 50% Campylobacteria, and >10% of Anaerolineae, Desulfobulbia, Dusulfuromonadia, Gracilibacteria, and Parcubacteria (Figure 4C).

Snap-Snap and Ultra-No-Chi-Chi Chimneys had similar distributions of microbial classes. Snap-Snap Chimney, the slightly cooler and shallower of the twoUrashima Chimneys, had a larger percentage of Campylobacteria and Deltaproteobacteria than Ultra-No-Chi-Chi Chimney with regards to ORFs (Figure 4A). By SSU rRNA genes of the composite assembly of Snap Snap had 25% Campylobacteria, 12% unclassified Bacteria, 12% Deltaproteobacteria, 10% Alphaproteobacteria, 10% Gammaproteobacteria, and less than 10% of Bacteroidetes, Flavobacteria, and unclassified Proteobacteria (Figure 4B). Amplicon sequencing showed 56% Campylobacteria, 10% Alphaproteobacteria, 10% Zetaproteobacteria, and less than 10% of Bacteroidia, Desulfobulbia, Gammaproteobacteria, and Gracilibacteria (Figure 4C). Ultra-No-Chi-Chi’s composite assembly showed the SSU genes to be 35% unclassified Bacteria, 30% Campylobacteria, 10% Alphaproteobacteria, 10% Gammaproteobacteria, and 15% unclassified Proteobacteria (Figure 4B). Amplicon sequencing showed 56% Campylobacteria, 8% Alphaproteobacteria, 8% Bacteroidia, 5% Gammaproteobacteria, 5% Gracilibacteria, and 18% Zetaproteobacteria (Figure 4C). Ochre Chimney’s ORFs showed a similar distribution to the Castle Chimney, with 26% unclassified Bacteria, 18% Gammaproteobacteria, 12% unclassified Proteobacteria, 21% Alphaproteobacteria, and 11% unclassified Chloroflexi (Figure 4A). Composite metagenome SSU genes showed 53% unclassified Bacteria, 12% Alphaproteobacteria, 16% Gammaproteobacteria, 10% Nitrososphaeria, and 9% unclassified Proteobacteria (Figure 4B). Amplicon sequencing showed 54% Gammaproteobacteria, 15% Alphaproteobacteria, and less than 10% of Bacteroidia, Dehalococcoidia, Parcubacteria, Phycisphaerae, and Planctomycetes (Figure 4C).

Pagoda Chimney had the highest alpha diversity of both Shannon and Simpson indices in contigs, while Ochre Chimney had the highest alpha diversity regarding amplicon-generated ASVs. (Supp. Table 2).

### Gene Abundances

The most abundant gene present in all chimneys was a putative transposase gene (Figure 5). Notably, an ammonium transporter gene was also present in the top 15 most abundant genes, with higher abundance at Castle Chimney and lower abundance at the Pagoda Chimney. DNA-directed RNA polymerase was present at all chimneys, with higher abundances in Ochre and Pagoda. Two chemotaxis genes were found at high abundance in the Snap-Snap Chimney but less abundant in the other chimneys.

**Figure 5.**
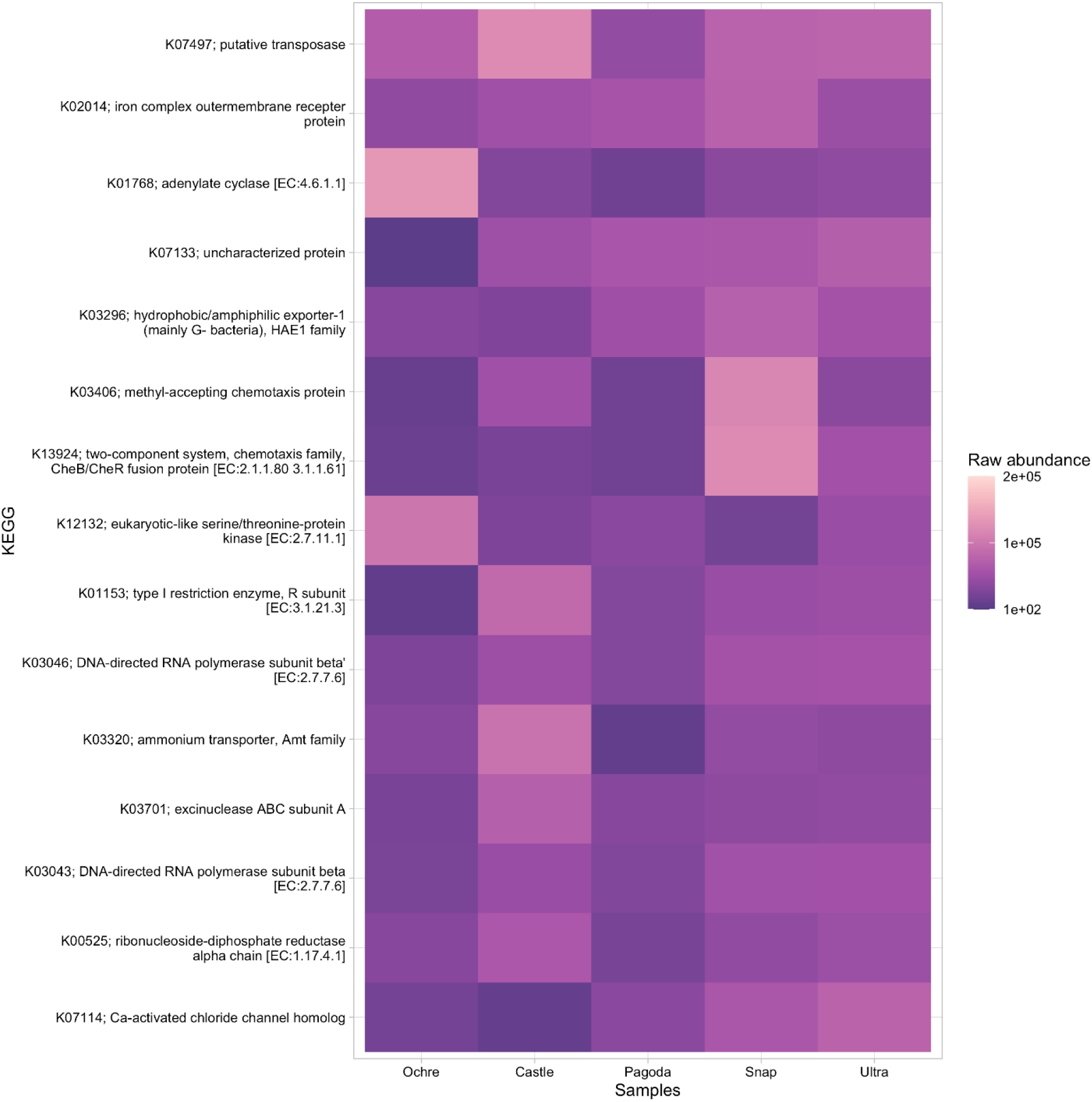
Heatmap of the top 15 most abundant KEGG genes found in all five chimneys. Presence is measured in raw abundance of reads.

### Metabolic potential

#### Carbon cycling

Chemosynthetic primary producers at hydrothermal vent chimneys fix carbon via the reverse tricarboxylic acid cycle (rTCA), the Calvin-Benson-Bassham (CBB) cycle, and the Wood-Ljungdahl (WL) pathway. The rTCA cycle uses the enzyme ATP citrate lyase (*aclB)* to take CO2 and water to make carbon compounds that can be used for energy by microorganisms in low-oxygen environments. The *aclB* gene was found to be present in all five chimneys (Figure 6). Castle and Ochre Chimneys both had Nitrospirae and Campylobacteria identified *aclB* genes (Supp. Tables 3 and 4). Pagoda Chimney, *aclB* was found in unclassified archaea and bacteria, as well as Thermoplasmata, Candidatus Bipolaricaulota, Chloroflexi, Aquificae, and Campylobacteria. At Snap-Snap and Ultra-No-Chi-Chi Chimneys, Campylobacteria *aclB* genes dominated in relative abundance (Supp. Tables 5-7).

**Figure 6.**
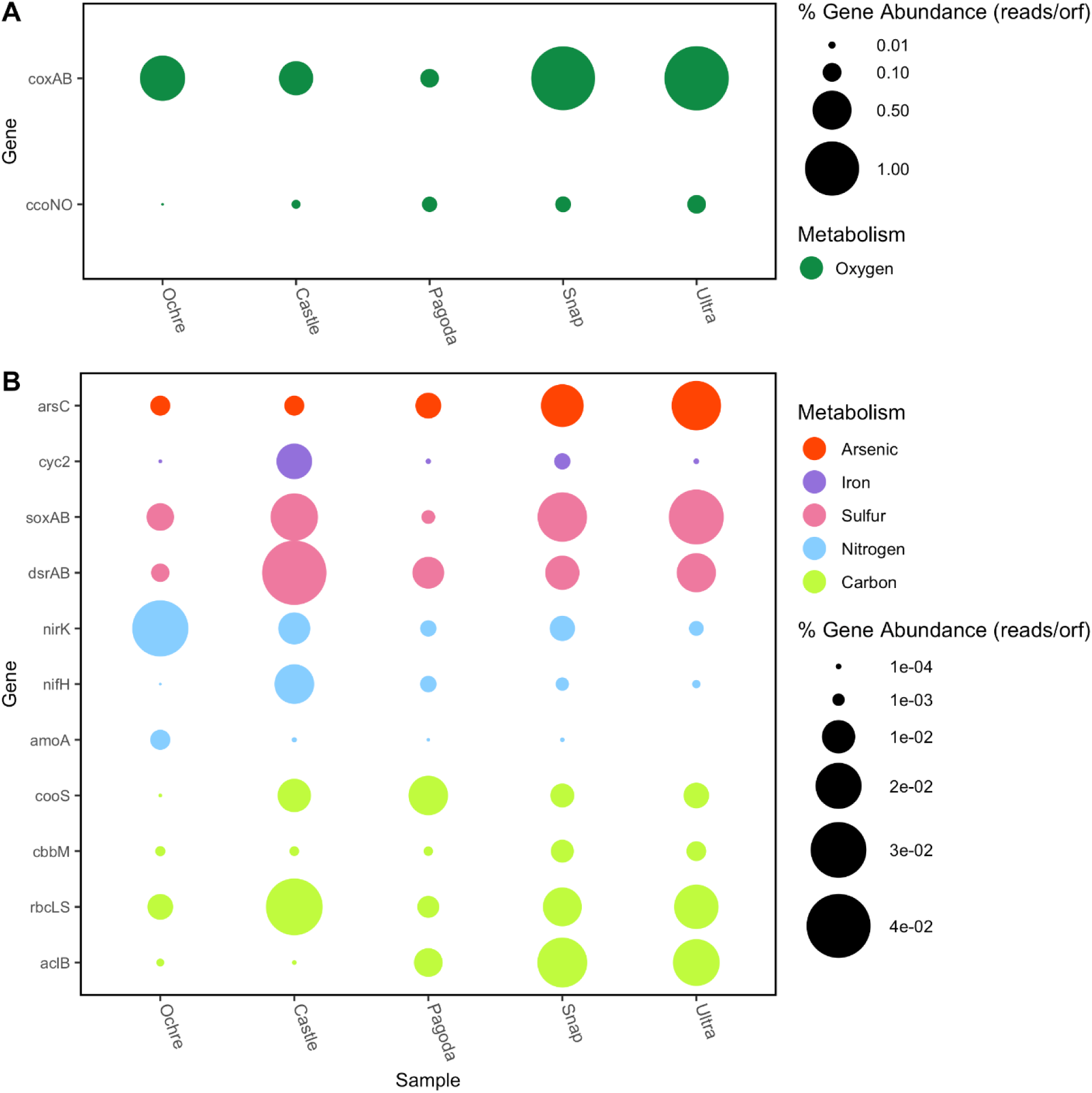
Bubble plot of the relative abundance of key metabolic genes found in each chimney. Relative abundance is measured in the number of reads per ORF divided by total number of reads. (A) Oxygen metabolism genes represented by dark green bubbles. (B) Arsenic, Iron, Sulfur, Nitrogen and Carbon genes. Red bubbles represent arsenic cycling genes, purple bubbles represent iron cycling genes, pink bubbles represent sulfur cycling genes, blue bubbles represent nitrogen cycling genes, and lime green bubbles represent carbon cycling genes.

The CBB cycle was identified using the RuBisCO protein, shown here by the presence of type I and type II RuBisCO genes, *rbcLS,* and *cbbM*, which have varying affinities for oxygen. These genes were found to be differentially abundant at all five chimneys, with *rbcLS* dominating (Figure 6). The *rbcLS* ORFs found were identified as Alphaproteobacteria, Chloroflexi, Proteobacteria, and Gammaproteobacteria across all five chimneys (Figure 6, Supp. Tables 3-7). The Castle and Ochre Chimneys showed similar assignments of *rbcLS*-assigned taxa, with Proteobacteria dominating and a low presence of archaeal *rbcLS* genes (Supp. Tables 3 and 4). In contrast, Pagoda Chimney and both Urashima Chimneys showed a higher presence of archaeal *rbcLS* genes (Table 2, Supp. Tables 5-7). The *cbbM* gene was identified as Alphaproteobacteria in every chimney and Gammaproteobacteria in every chimney but Pagoda, which was dominated by Methanomicrobia and Thermococci. The two Urashima chimneys had a large abundance of Zetaproteobacterial *cbbM* genes (Supp. Tables 6 and 7).

The Wood-Ljungdahl pathway reduces CO2 to carbon monoxide using the enzyme carbon monoxide dehydrogenase (*cooS* and *acsA,* both represented here by *cooS*) and then creates acetyl-CoA using acetyl-CoA synthase (25). The Wood-Ljungdhal pathway was found to be present across all five chimneys (Figure 6). This gene was least abundant at Ochre Chimney and most abundant at Pagoda Chimney (Figure 6). Deltaproteobacteria, Gammaproteobacteria, and Nitrospirae identified *cooS* all were found at higher relative abundances at Castle Chimney, Chloroflexi, Deltaproteobacteria, and Methanopyri *cooS* were the most abundant at Pagoda Chimney, and Deltaproteobacteria and Nitrospirae *cooS* were found to be in the highest relative abundance at the two Urashima chimneys (Supp. Tables 3-7).

#### Nitrogen cycling

Three nitrogen cycling genes were annotated and taxonomically assigned for each chimney: methane/ammonia monooxygenase subunit A (*amoA*), nitrogenase iron protein (*nifH*), and nitrate reductase (*nirK*), a gene responsible for the denitrification of nitrite to nitric oxide (NO). The least abundant nitrogen cycling gene across all five chimneys was *amoA* (Supp. Tables 3-7). Nitrification as represented by the *amoA* gene showed the greatest representation in the Ochre Chimney, with corresponding taxa including Betaproteobacteria, Deltaproteobacteria, Gammaproteobacteria, and Nitrososphaeria, unclassified Thaumarchaeota and unclassified Archaea (Supp. Table 3). Interestingly, *amoA* is completely absent from the Ultra-No-Chi-Chi Chimney while this gene was identified at Snap-Snap, its geographic neighbor.

Pagoda Chimney had the lowest total abundance of nitrogen metabolism genes. dinitrogen fixation by *nifH* was dominated by Methanopyri, Nitrospirae, Firmicutes, Gammaproteobacteria, and Deltaproteobacteria (Figure 6, Supp. Table 5). Most of the ORFs assigned to *nifH* genes were found to be associated with Methanopyri, a class of methanogenic Euryarchaeota that are likely nitrogen fixers. The Castle Chimney had a large proportion of *nifH* genes dominated by Proteobacteria, while the two Urashima chimneys had small abundances of *nifH* genes identified as Deltaproteobacteria, Thermodesulfobacteria, and Archaeoglobi (Supp. Tables 3, 6, and 7).

The most prevalent form of nitrogen metabolism across the samples except the Pagoda and Castle Chimneys was denitrification via *nirK* (Figure 6). Each chimney had a unique taxonomic distribution of *nirK* taxa, with high relative abundances in Bacteroidetes, Gammaproteobacteria, and unclassified Bacteria. Ochre Chimney had the highest diversity of *nirK* genes across both archaea and bacteria, with most of the reads assigned to Nitrososphaeria (Supp. Tables 3-7).

#### Sulfur cycling

Two sulfur cycling genes were examined in all five chimneys: dissimilatory sulfite reductase *dsrAB*, and thiosulfotransferase *soxAB*. Across all five chimneys, *dsrAB* was found present in Alphaproteobacteria, Deltaproteobacteria, Gammaproteobacteria, Proteobacteria, and unclassified Bacteria. It was also found in the four active chimneys as Acidobacteria and Nitrospirae. The *soxAB* genes were represented across similar taxa as *dsrAB* genes in all five chimneys. At Ochre Chimney, the Gammaproteobacterial and Alphaproteobacterial *dsrAB* genes found present are the oxidative version of *dsrAB,* indicating that these organisms are sulfur-oxidizing bacteria. Alphaproteobacteria, Gammaproteobacteria, and unclassified Bacteria *dsrAB* genes were present in all chimneys and Deltaproteobacteria and Campylobacteria were present in the four active chimneys (Supp. Tables 3-7).

#### Iron cycling

To observe iron metabolisms in the chimneys, the iron oxidase gene *cyc2* was examined since it is useful as an indicator of microbial iron oxidation. The presence of iron was confirmed at the Castle Chimney and two Urashima chimneys (Table 1). However, *cyc2* genes were found present in both the Pagoda Chimney and the Ochre Chimney. The most abundant *cyc2* gene taxon was Gammaproteobacteria, found in all chimneys except for Ultra-No-Chi-Chi (Supp. Tables 3-7). At Snap-Snap Chimney Alphaproteobacteria, Gammaproteobacteria, Bacteroidetes, and unclassified Bacteria had the *cyc2* gene. In contrast, Ultra-No-Chi-Chi only had ORFs assigned to Proteobacteria and Aquificae *cyc2* genes (Supp. Tables 3-7). In FeGenie, *cyc2* wasn’t identified (Supp. Fig. 1). The iron complex outer membrane receptor gene, involved in the acquisition and uptake of iron, was found to be present across all five chimneys, with the largest relative abundance found in the Snap-Snap Chimney (Figure 5).

**Table 1.**
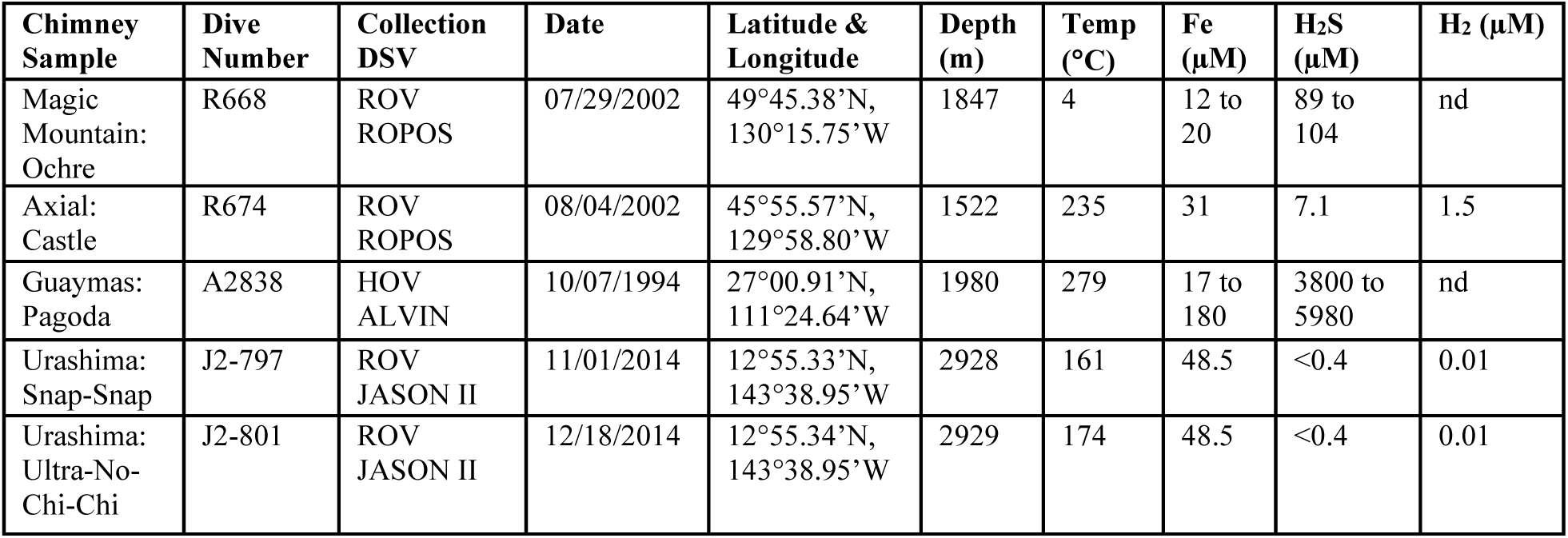
A summary of samples collected from five different hydrothermal vent chimneys.

| Chimney Sample | Dive Number | Collection DSV | Date | Latitude & Longitude | Depth (m) | Temp (°C) | Fe (μM) | H<sub>2</sub>S (μM) | H<sub>2</sub> (μM) |
| --- | --- | --- | --- | --- | --- | --- | --- | --- | --- |
| Magic Mountain: Ochre | R668 | ROV ROPOS | 07/29/2002 | 49°45.38'N, 130°15.75'W | 1847 | 4 | 12 to 20 | 89 to 104 | nd |
| Axial: Castle | R674 | ROV ROPOS | 08/04/2002 | 45°55.57'N, 129°58.80'W | 1522 | 235 | 31 | 7.1 | 1.5 |
| Guaymas: Pagoda | A2838 | HOV ALVIN | 10/07/1994 | 27°00.91'N, 111°24.64'W | 1980 | 279 | 17 to 180 | 3800 to 5980 | nd |
| Urashima: Snap-Snap | J2-797 | ROV JASON II | 11/01/2014 | 12°55.33'N, 143°38.95'W | 2928 | 161 | 48.5 | <0.4 | 0.01 |
| Urashima: Ultra-No-Chi-Chi | J2-801 | ROV JASON II | 12/18/2014 | 12°55.34'N, 143°38.95'W | 2929 | 174 | 48.5 | <0.4 | 0.01 |

#### Arsenic metabolism

Arsenate reductase, *arsC,* allows for the reduction of arsenate and each of the five chimneys contained this gene in at least 12 different taxa including Gammaproteobacteria, Bacteroidetes, Proteobacteria, and unclassified Bacteria (Supp. Tables 3-7). Ultra-No-Chi-Chi Chimney had the most *arsC* genes present. In the Castle and Ochre Chimneys, Gammaproteobacteria had the most ORFs assigned, while Campylobacteria dominated the two Urashima chimneys. The Pagoda Chimney had the largest abundance of Bacteroidetes identified *arsC* genes (Supp. Tables 3-7).

#### Oxygen metabolism

The relative abundance of two cytochrome c oxidase genes (*ccoNO* and *coxAB*) was evaluated as an indicator of aerobic respiration. The *ccoNO* gene was more abundant at the Ultra-No-Chi-Chi, Snap-Spap, and Pagoda Chimneys, with the least amount detected at the Castle and Ochre Chimneys (Figure 6). At Ochre Chimney, Bacteriodetes, Flavobacteriia, and Gemmatimondetes had the highest relative abundance of ORFs assigned to *ccoNO* genes (Supp. Table 2). Castle Chimney had a large abundance of Bacteroidetes and Gammaproteobacteria *ccoNO* genes (Supp. Table 4). Chlorobi, Flavobacteriia, Bacteriodetes, and Cytophagia *ccoNO* genes were most abundant at Pagoda Chimney (Supp. Table 4). At Snap-Snap; Acidobacteria, Bacteriodetes, Chlorobi, and Flavobacteriia had the most *ccoNO* genes assigned (Supp. Table 6). Ultra-No-Chi-Chi had a similar abundance of *ccoNO* genes, with the addition of Deltaproteobacteria having a large relative abundance (Supp. Table 7). Overall, the *coxAB* gene was much higher than the *ccoNO* genes at all chimneys except Pagoda, which had the lowest relative abundance. There were large relative abundances of *coxAB* genes assigned in Alpha and Gammaproteobacteria in all the chimneys, and Pagoda and Ultra-No-Chi-Chi chimneys had a large relative abundance of Campylobacteria assigned to *coxAB* genes (Supp. Table 3-7).

## DISCUSSION

The analysis of the metagenomes of five hydrothermal vent chimneys around the Pacific Ocean demonstrates a set of functional core genes that all locations share while also elucidating the differences in community composition and metabolic potential. After examining the binned MAGs in each chimney, the SSU rRNA taxonomic community compositions, and some common genes for carbon fixation, nitrogen, sulfur, iron, arsenic, and oxygen metabolism, each of the chimneys were found to have a distinct metabolic profile.

PERMANOVA analysis showed no statistical significance to the chimneys clustering based on geographic location, activity, depth, or temperature; however, this is likely due to having no sample replicates. Despite a lack of statistically significant data, each chimney site has a unique community of microbes; however, there were noticeable trends. Similarities between Snap-Snap and Ultra-No-Chi-Chi Chimneys are likely due to them being close in proximity and with similar chemical profiles. Similarities in metabolic potential and high prevalence of Gammaproteobacteria between Castle Chimney and Ochre Chimney likely are due to the shallow depth of Castle Chimney and the inactivity of Ochre Chimney allowing for growth of more aerobic microorganisms. This is supported by the high relative abundance of the *coxAB*, indicating aerobic respiration is occurring in the presence of a high concentration of oxygen (26).

The relative abundance of transposase genes in all five chimneys indicates horizontal gene transfer as a likely method of adaptation to the extreme environment of the chimneys (27, 28). Horizontal gene transfer among microbial taxa increases the phenotypic diversity of the chimney communities for microbes to better respond and adapt to environmental gradients (27). This is evident when examining extremophiles that commonly reside at hydrothermal vents. The extremophile *Fervidobacterium* showed transposases as indicators of horizontal gene transfer are common in thermophilic microbes (29).

The presence of an ammonia transport gene supports the high instance of ammonia oxidation at each of these chimneys. The ubiquity of ammonia transport genes across the chimneys suggests that microbes are accessing environmental nitrogen for assimilative or dissimilative processes (30).

Arsenic is inferred to be present at all five chimneys based on the ubiquity of the *arsC* gene. A diverse number of microbes can metabolize arsenic using *arsC* (31). Since nine different classes of microorganisms at these chimneys have an arsenic reductase gene, arsenic is likely present and detoxification is necessary for survival.

### Ochre Chimney

Ochre Chimney is an inactive, weathered chimney; therefore, the microbial community must gain its energy from the metabolism of solid minerals rather than the reduced chemicals present in the venting fluid of an active chimney. The lack of venting fluid at inactive chimneys allows for more stable metabolic activity and cooler temperatures (32). At Ochre Chimney, this is supported by the lower abundance of Type I restriction enzymes and genes for chemotaxis. The lower relative abundance of these genes suggests less demand for microbes to respond quickly to changing chemical gradients since the sources of energy in the chimney sediment are relatively stable (33).

The abundance of *rbcLS* and *cbbM* assigned taxa at Ochre Chimney confirms carbon cycling at inactive chimneys is done via the CBB cycle (14). Other studies examining the metabolic potential of inactive hydrothermal vent chimneys on the East Pacific Rise have identified the CBB cycle in autotrophic Gammaproteobacteria (34). Notably, Ochre also has two archaeal classes assigned to *rbcLS*, one in the Thaumarchaeota phylum and one at the unclassified Archaeal level. Archaeal RuBisCO genes are putatively involved in carbon dioxide fixation or AMP and nucleotide scavenging pathways (35). Thaumarchaeota, specifically Nitrososphaeria, are known to be common in inactive chimneys and can metabolize low concentrations of nitrogen and carbon (7).

As *nirK* is largely used in archaea for ammonia oxidation and in bacteria for denitrification of nitrite, the relatively high number of different taxa that have a *nirK* gene for denitrification indicates an abundance of nitrite as an electron donor for lithotrophic growth (36). At the hydrothermal vents of Explorer Ridge, nitrate was more prevalent than nitrite, which explained the high relative abundance of *nirK* (37). The relatively large abundance of *amoA* genes could be due to increased ammonium found at the inactive chimney due to the breakdown of organic matter (38). Both bacterial and archaeal *amoA* were identified and are likely critical in the nitrification process at Ochre Chimney.

The availability of sulfur is a distinguishing factor in community composition between inactive and active chimneys (7). Since there is no data on the hydrogen sulfide concentrations at Ochre Chimney, the presence of different sulfur compounds can only be inferred by the differential abundance genes for sulfur metabolism. Sulfate reduction by Deltaproteobacteria dominates at inactive chimneys, which in turn can change the mineral composition of the chimney with the production of pyrite (7). The taxonomic identification overlap with *dsrAB* and *soxAB* indicates that if dissimilatory sulfate reduction is occurring, thiosulfate oxidation could be occurring concurrently in the same taxa. This type of functional redundancy has been shown to increase ecological stability and resilience to disturbance, like the inactivation of a chimney (39).

The relatively low abundance of Gammaproteobacterial *cyc2* genes at Ochre Chimney indicates that iron oxidation may not be as prevalent at inactive, weathered chimney structures. Gammaproteobacteria are primary colonizers of inactive chimneys, as they can oxidize sulfur present in the chimney structure (34). These Gammaproteobacteria may act as a catalyst in the weathering of inactive iron-sulfide chimneys, which could indicate that the Ochre Chimney was towards the end of the weathering process (14).

Gammaproteobacteria tend to favor environments with higher oxygen (32). The *ccoNO* gene encodes a cbb3-type cytochrome c oxidase subunit I/II, which has a high affinity for oxygen and is more prevalent in lower oxygen concentrations, while the *coxAB* gene encodes an aa3-type cytochrome c oxidase subunit I/II, which has a low affinity for oxygen and is more prevalent in higher oxygen concentrations (40). Ochre Chimney has a large relative abundance of Gammaproteobacteria and a very small relative abundance of *ccoNO* cytochrome c oxidase genes, further supporting that oxygen concentration is higher at this chimney.

### Castle Chimney

The main pathway for carbon fixation at Castle Chimney is via the CBB cycle, with most *rbcLS* genes associated with different Proteobacterial classes. By metagenomics of Axial Seamount chimney, Gammaproteobacteria were found to be the largest contributing taxon to the CBB cycle (18). Gammaproteobacteria tend to favor environments with higher oxygen and lower concentrations of sulfide (41). The higher concentration of oxygen present at Castle Chimney may account for the high relative abundance of Gammaproteobacterial *rbcL* genes. The presumed high concentration of oxygen is also evidenced by the relatively low abundance of *ccoNO* genes.

Despite the abundance of *rbcLS*, Campylobacteria and Nitrospirae fix carbon via the rTCA cycle at Castle Chimney, which has been previously demonstrated at Axial Seamount (18). rTCA is favored over CBB in oxygen-limited environments (42), which could explain the higher relative abundance of Gammaproteobacterial CBB cycling genes over Campybacterial rTCA cycling genes.

The dominance of denitrification as evidenced by the large proportion of bacterial and archaeal *nirK* genes is supported by previous analyses, which classified *nirK* transcripts to Thaumarchaeota and other ammonia-oxidizing archaea at Axial Seamount (18). Organisms that had the ammonia oxidation gene, *amoA,* always had *nirK* genes as well, indicating that these pathways are likely co-occurring in ammonia-oxidizing archaea which is hypothesized to be due to the decentralization of gene expression to maintain genetic diversity in variable environments like hydrothermal vent chimneys (43).

Since Castle Chimney likely has anaerobic sulfide-oxidizing bacteria since nitrite reduction genes and sulfur oxidation genes were identified as Gammaproteobacteria. As with Gammaproteobacterial sulfide oxidation via *dsrAB*, some Alphaproteobacteria couple denitrification via *nirK* with thiosulfate oxidation via *soxAB* in deep subsurface environments (44). Since many of these metabolic pathways have been shown to co-occur, it demonstrates that the microbes present in Castle Chimney likely may be capable of gaining electrons from several different sources, as evidenced by the co-occurrence of Alphaproteobacterial *nirK* and *soxAB*.

Castle Chimney iron oxidation is dominated by Gammaproteobacteria, a class that has been previously identified on the Juan de Fuca Ridge (15). Zetaproteobacteria, an iron-oxidizing bacteria commonly found at hydrothermal vents, is notably absent in Castle Chimney. This could be due to misidentification, or the higher temperature and lower abundance of iron at Castle could influence a higher proportion of Gammaproteobacterial iron oxidation, as Zetaproteobacteria tend to prefer lower temperatures (45).

### Pagoda Chimney

Shaped like a mushroom or a Pagoda topped with a domed cap and many flanges coming out of the trunk, Pagoda Chimney’s vent fluid is channelized through the flanges and up and over its cap, collecting in the center and creating several microenvironments of differing temperatures and chemistries (19). These different habitats introduce a need for microbes to adapt quickly to an ever-changing environment which is supported by the enrichment of transposase genes and therefore an increased potential for horizontal gene transfer (46). Pagoda Chimney has the most taxonomic diversity of *rbcLS* genes, with a higher presence of archaeal *rbcLS* genes, likely due to the physical structure of the chimney allowing for many temperatures and chemical gradients (47). It has been shown that RuBisCO can also be used for nucleotide salvage rather than carbon fixation in archaea, which could explain the high abundance of archaeal *rbcLS* genes present (48).

Guaymas Basin is characterized by organic-rich sediment and high phytoplankton productivity supporting heterotrophic metabolisms (49). These organic-rich sediments lead to a large amount of hydrogen to be used for energy by organisms like Methanopyri (20). Sulfate reduction is common among microbial communities at Guaymas Basin hydrothermal chimneys. Sulfate-reducing bacteria degrade the plentiful hydrocarbons found at this site which in turn create H2 that can be used by methanogens in low oxygen environments (46). Both the oxidative and reductive versions of *dsrAB* were present in Pagoda. The only instance of archaeal *dsrAB* genes present were in the Archaeoglobi class, the only known archaeal class that is hyperthermophilic with a sulfate-reducing metabolism (50). As expected, Methanopyri was abundant in Pagoda Chimney and likely utilizes Wood-Ljundhal pathway for carbon fixation and methanogenesis (51). Multiple hydrogenase uptake genes were found present at Pagoda, to further supporting *Methanopyri* methanogenesis. There is likely deoxygenation occurring inside Pagoda chimney as supported by low abundance of *coxAB* and *ccoNO* (19). This deoxygenation creates environments favorable for anaerobes such as *Methanopyri*.

### Snap-Snap Chimney

Back arcs, like the Urashima Vent Field, can have a wide variation in pH, dissolved gases, and metal concentrations due to variations in the magma chemistry (22). The high abundance of genes for chemotaxis at Snap-Snap indicates a steep gradient of reduced chemicals needed for growth. The presence of *napA*, periplasmic nitrate reductase, is indicative of low oxygen presence and anaerobic respiration (52). The low oxygen concentration is evidenced at Snap-Snap by the large relative abundance of *ccoNO* cytochrome c oxidase genes.

Snap-Snap Chimney had a high abundance of Gammaproteobacterial *aclB* and *rbcLS* genes and the most *cbbM* genes out of all the chimneys, consistent with previous analyses of the CBB cycle on the Mariana back-arc (22). The presence of the *rbcLS* gene mapped to *Deinococci*, an extremophile chemoorganotroph, and the high abundance of chemotaxis proteins could further indicate that the Snap-Snap Chimney is an extreme environment with variable chemical, temperature, and nutrient gradients.

Based on the relative abundances of dissimilatory nitrogen metabolism genes found at Snap-Snap Chimney, a large amount of nitrite is likely used as an energy source. Previous analyses of archaeal denitrifiers have found that accumulation of organic material can increase *nirK* gene abundances, indicating that there may be organic material build-up at Snap-Snap Chimney as there are archaeal *nirK* genes present (53).

At Snap-Snap Chimney, there is a higher abundance of Campylobacteria, indicating a high prevalence of sulfide in the vent fluid (22, 54). Deltaproteobacteria and Gammaproteobacteria have been found to have the ability to couple sulfate reduction with sulfur oxidation using the oxidative form of *dsrAB* (55). Since these taxa both have both sulfur genes, coupled sulfur oxidation with sulfate reduction is likely occurring.

Snap-Snap Chimney is characterized by high concentrations of iron (22), which is hypothesized to be due to low pH from magmatic volatiles on the Mariana back-arc (56). Snap-Snap Chimney has several taxa with to the *cyc2* gene; however, no Zetaproteobacterial *cyc2* genes were identified in contrast to previous research (23). This could be because as it is a relatively new class, the Zetaproteobacterial *cyc2* ORFs are folded into the Gammaproteobacterial or unclassified bacterial category during annotation (57).

### Ultra-No-Chi-Chi Chimney

Notably, Ultra-No-Chi-Chi has nearly the same number of *aclB* and *rbcLS* genes, demonstrating that both the rTCA and CBB pathways are used for carbon fixation. The rTCA cycle is likely performed by Campylobacteria, while the CBB cycle has a higher diversity of taxa. The highly relative abundance of both *ccoNO* and *coxAB* genes indicates that Ultra-No-Chi-Chi has a broad oxygen gradient, allowing for anaerobic and aerobic organisms to fix carbon.

Unexpectedly, Ultra-No-Chi-Chi has a lower abundance of genes coding for iron receptors and *cyc2,* which could indicate that there is less iron present. This was unexpected, as other Urashima chimneys have been characterized as iron-dominated (23). However, Zetaproteobacterial *nirK* and SSU genes were found. Iron oxidation is coupled with denitrification via *nirK* in Zetaproteobacteria (58). Therefore, the presence of Zetaproteobacterial *nirK* and SSU genes confirms their presence and therefore iron oxidation at this chimney despite no *cyc2* genes mapping to that class.

Metagenomic analysis of these five hydrothermal vent chimneys demonstrates how the chemical composition of the chimney impacts the microbes that reside there and their potential metabolisms. Despite a unique collection of microorganisms, there were several uniting characteristics. Genes for DNA repair, chemotaxis, and transposases were present at higher abundances at hydrothermal vent chimneys compared to other environmental microbial communities and could be a uniting identifier for these communities to adapt to the ever-changing chemical and physical conditions. Gammaproteobacteria dominated Ochre and Castle Chimney while Campylobacteria were more prevalent at Pagoda, Snap Snap, and Ultra-No-Chi-Chi. The relative abundances of oxygen and carbon metabolism genes at each of the chimneys tell a distinct story of the availability of these compounds as energy sources in both active and inactive chimneys. High oxygen metabolism genes coupled with low carbon fixation genes could be used as a unique identifier for inactive chimneys, as shown with Ochre Chimney. The differences in carbon fixation genes and their metabolic plasticity demonstrate that chimney microbes can adapt to varying chemical compositions of the chimneys and that many of these metabolic pathways tend to be functionally redundant to thrive in a dynamic ecosystem.

## METHODS

### Study Sites and Sampling

Chimneys were collected from Axial Seamount, Magic Mountain, Guaymas Basin, and two chimneys from the Urashima Vent Field along the Mariana back-arc using either a self-sealing scoop sampler or biobox. At Axial Seamount, Castle Chimney was collected on August 4^th^, 2002 with ROV ROPOS aboard the R/V Thomas G. Thompson. The Ochre Chimney at Magic Mountain Vent Field was collected on July 29^th^, 2002 with ROV ROPOS aboard the R/V Thomas G. Thompson. The Pagoda Chimney at the Guaymas Basin Vent Field was collected on October 7^th^, 1994 with HOV ALVIN aboard the R/V Atlantis. Two chimney samples, Snap-Snap and Ultra-No-Chi-Chi, were collected with ROV Jason aboard the R/V Roger Revelle from the Urashima Vent Field. Snap-Snap Chimney was sampled on November 1^st^, 2014 and Ultra-No-Chi-Chi Chimney was sampled on December 18^th^, 2014. All chimney samples from all sites were immediately preserved with RNAlater, and then stored at -80°C until DNA could be extracted. Geochemistry data that has been previously published (59, 60).

### Amplicon DNA Extractions, Sequencing, and Sequence Processing

Genomic DNA was extracted from cell pellets using the Fast DNA SPIN Kit for Soil (MP Biomedicals, Santa Ana, CA) as previously published (61). Cell lysis was optimized using two rounds of bead beating for 45 sec at a power setting of 5.5 using the FastPrep instrument (MP Biomedicals) with samples being placed on ice between runs. Extracted DNA was quantified by a Qubit 2.0 fluorometer using high-sensitivity reagents (ThermoFisher Scientific, Waltham, MA).

The V3-V4 regions of the SSU rRNA gene were amplified via PCR-purified DNA using bacterial primers 340F and 784R (61, 62). The resulting amplicons were sequenced using a MiSeq (Illumina, San Diego, CA) as per the manufacturer’s protocol generating 2 x 300bp paired-end reads. The resulting reads were trimmed of primers using CutAdapt (63). The trimmed reads were then processed using the Divisive Amplicon Denoising Algorithm 2 (DADA2) v1.26.0 with pseudopooling following the previously described protocols (64–66) with R version 4.2.3 and using the Silva v138 database for assigning taxonomy. Further analysis was completed using phyloseq version 1.32 (67) and microbiome version 1.20 (68).

### Metagenomic Sequencing

The extracted DNA was separated to strands greater than 1 kb using an Aurora (Boreal Genomics, Vancouver, BC). Nextera indices were added to purified DNA fragments per the manufacturer’s protocol (Illumina, San Diego, CA). The indexed fragments were purified using AMPure XP Beads according to the manufacturer’s protocol (Beckman Coulter, IN). The library was quantified with a Qubit 2.0 fluorometer (ThermoFisher, MA) and sequenced on the Illumina MiSeq sequencer with v3.0 chemistry to generate 2×300 bp paired-end reads at Shannon Point Marine Center, WWU.

### Metagenomic Sequence Analysis

#### Assembly

After sequencing, the reads were assessed and trimmed for quality control using the program Trimmomatic v.0.40 (69) and were assessed for quality with FastQC v.0.11.9 (70).

All metagenomic analyses were done under the metagenomic pipeline, SqueezeMeta v.1.5.1 (71) under co-assembly mode using default parameters. Once trimmed for quality, the reads were assembled into contigs using MegaHIT with a minimum contig length of 200 base pairs (72). The assembled contigs were checked for rough taxonomy using MetaQuast (73). Genes were predicted using Prodigal (74). Small subunit (SSU) rRNA genes were pulled from contigs using Barrnap (75) and classified using the RDP naïve Bayesian classifier (76).

#### Functional annotation and taxonomic assignment

Gene sequences were compared for homology using Diamond v.2.7.14 (77). The genes were then functionally and taxonomically assigned using SqueezeMeta. Bowtie2 v.2.4.5 was used to estimate coverage and abundance (78). Gene abundances were calculated using STAMP (79), then the percent relative abundance of each gene was calculated by taking the raw read counts for each ORF, dividing it by the total reads in the sample, and multiplying by 100. Coverage values (bases mapped/ORF length) and normalized RPKM values were calculated using custom SqueezeMeta pipeline scripts. All outputs of taxonomic names present in this analysis reflect the names in the databases at the time of assembly and assignment. Contig assemblies were loaded into FeGenie to evaluate the abundance of different iron genes in each chimney (80). Outputs were visualized in R with the package ggplot2 v.3.3.5 (81).

#### Binning

After assembly, the contigs were binned into metagenomic assembled genomes (MAGs) using Metabat2 v.2.15 (82) and Maxbin2 v.2.2.7 (83). The outputs of both binning programs were merged using DAStool v.1.1.3 (84). Bins were checked for completeness and contamination using the program CheckM v.1.1.3 (85), then taxonomically assigned using the same LCA algorithm in SqueezeMeta as the taxonomic assignment of the genes.

#### Data analysis

In R v.3.6.3, the results of the SqueezeMeta co-assembly pipeline were imported using the R package SQMtools v.0.7.0 (86). Vegan (87) was used to determine permutational multivariate analysis of variance (PERMANOVA) among the chimneys with 999 permutations and Bray-Curtis distance method, non-metric multidimensional scaling (NMDS) based on Bray-Curtis distance of KEGG functions, and relative abundance and diversity of the taxa found at each chimney. The similarity of taxa was analyzed using Bray Curtis distance matrix and visualized with a dendrogram.

### Data Availability

The amplicon and shotgun sequencing data sets generated for this study have been deposited in the NCBI Sequence Read Archive under the BioProject accession no. PRJNA996601.

## Acknowledgments

This work was funded in part by the Fouts Foundation for the Enhancement of Student Research Experiences, and by the National Science Foundation, award OCE 1155756 (to CM).

---

**Supplemental Table 1.**
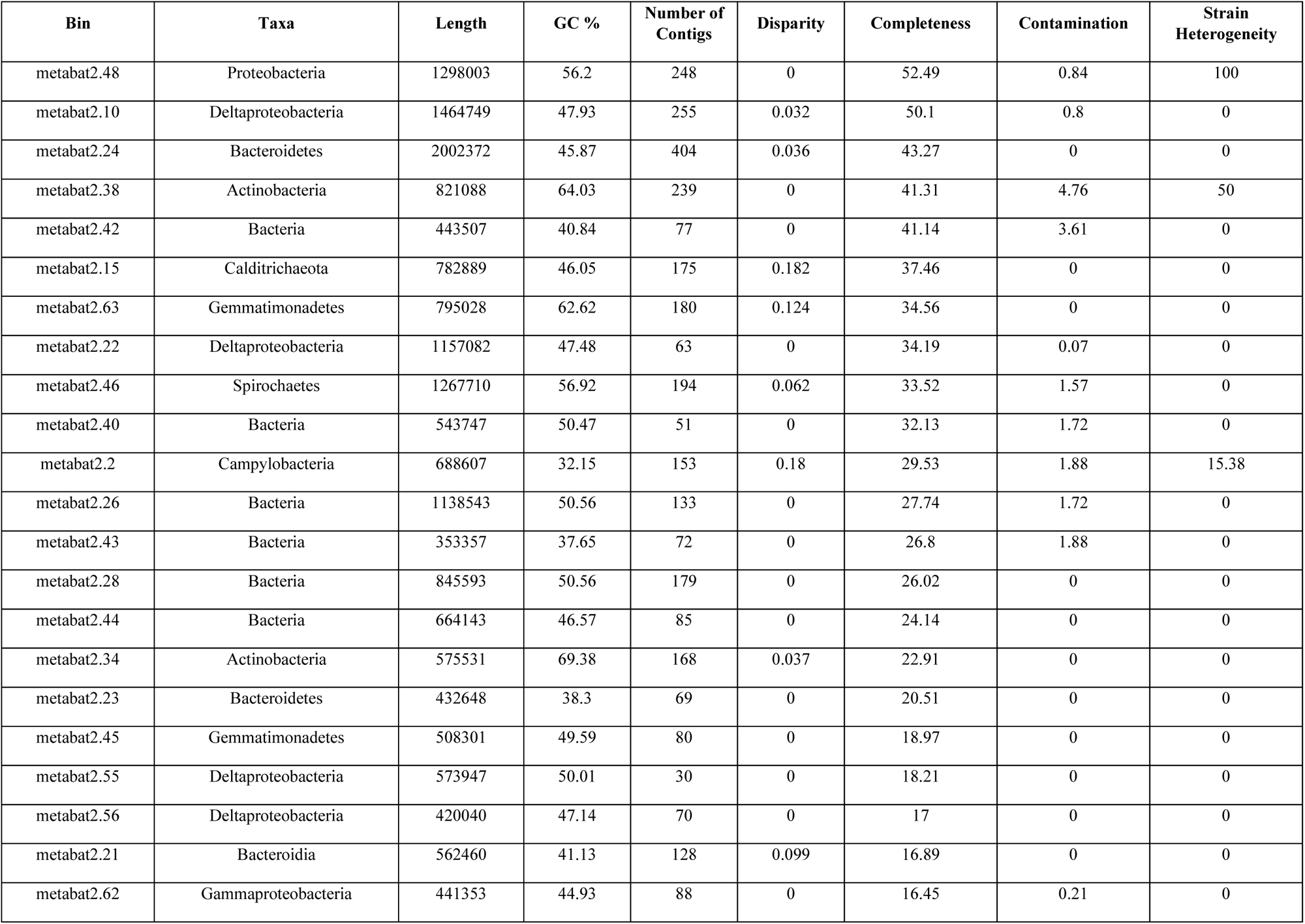
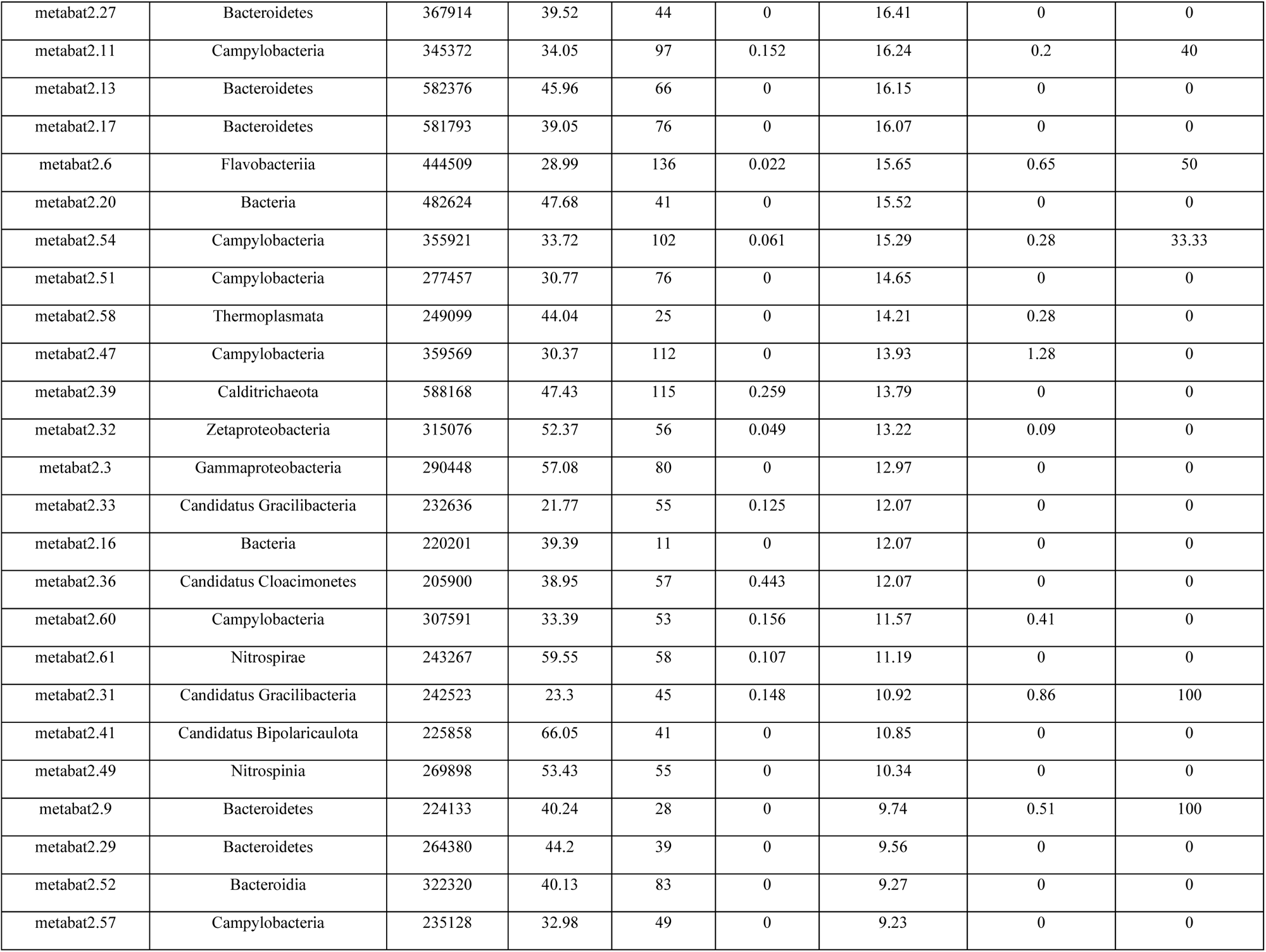
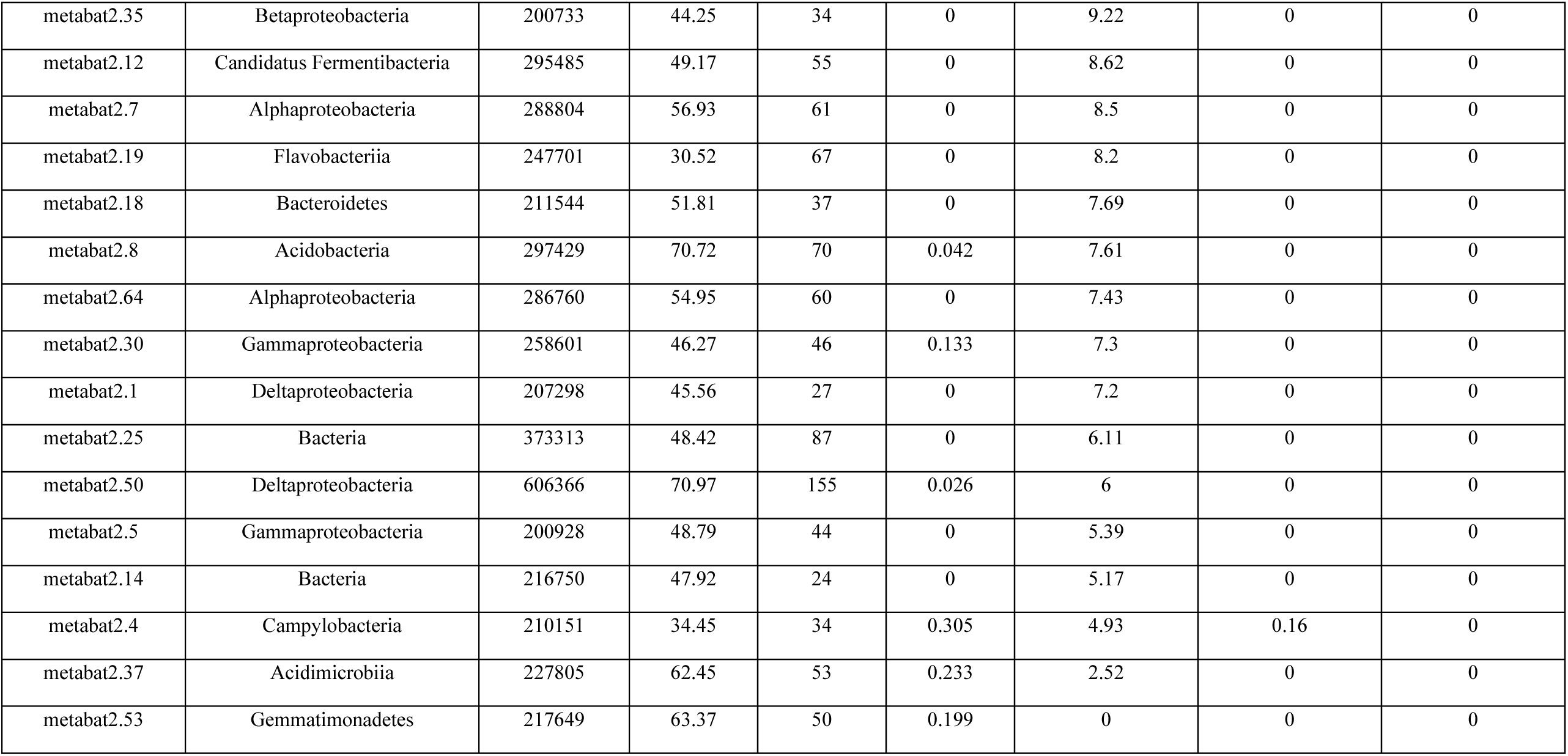
MAGs and their taxonomic assignments with completeness and contamination.

**Supplemental Table 2.** The Simpson and Shannon alpha diversity indices for the ASVs and Contigs from each Chimney.

| **Chimney** | **ASVs**<br>**Simpson** | **Contig**<br>**Simpson** | **ASVs**<br>**Shannon** | **Contig**<br>**Shannon** |
| --- | --- | --- | --- | --- |
| Ochre | 0.997 | 0.750 | 6.928 | 2.161 |
| Castle | 0.954 | 0.830 | 4.428 | 2.131 |
| Pagoda | 0.989 | 0.890 | 5.466 | 2.818 |
| Snap | 0.976 | 0.800 | 4.997 | 2.143 |
| Ultra | 0.937 | 0.860 | 4.194 | 2.510 |

**Supplemental Table 3.**
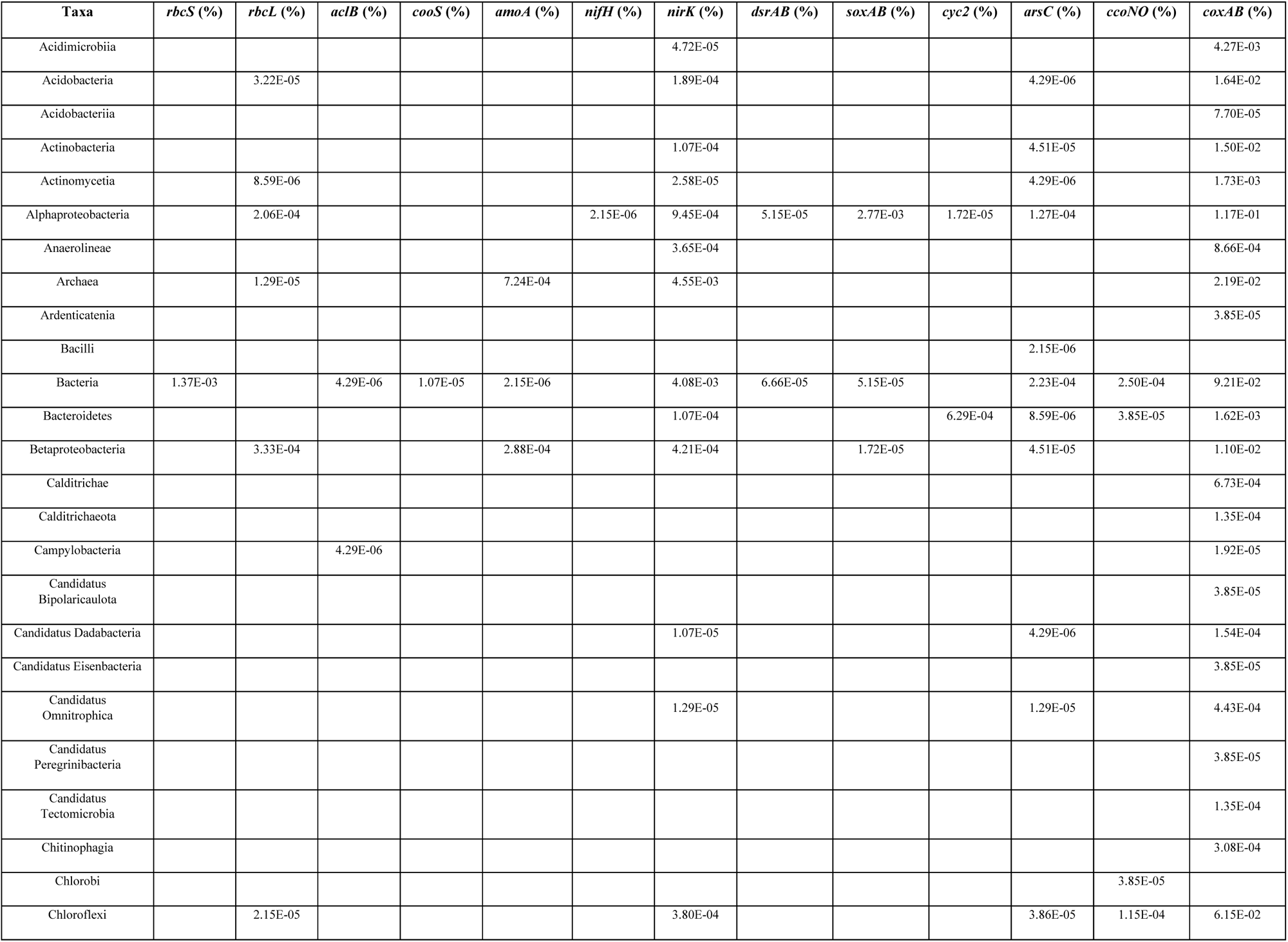
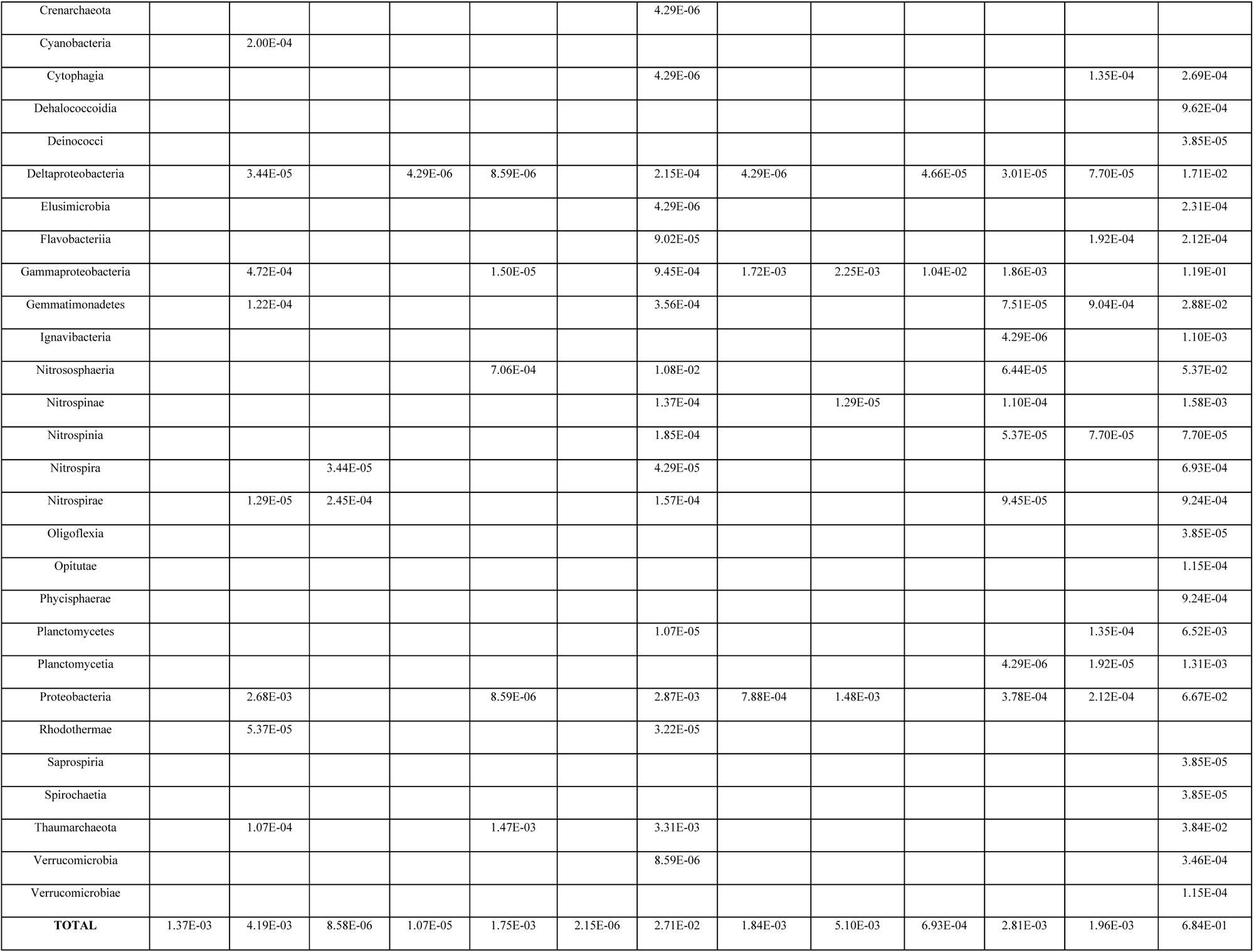
Relative abundance of different autotrophy genes and their taxonomic assignment for Ochre Chimney.

**Supplemental Table 4.**
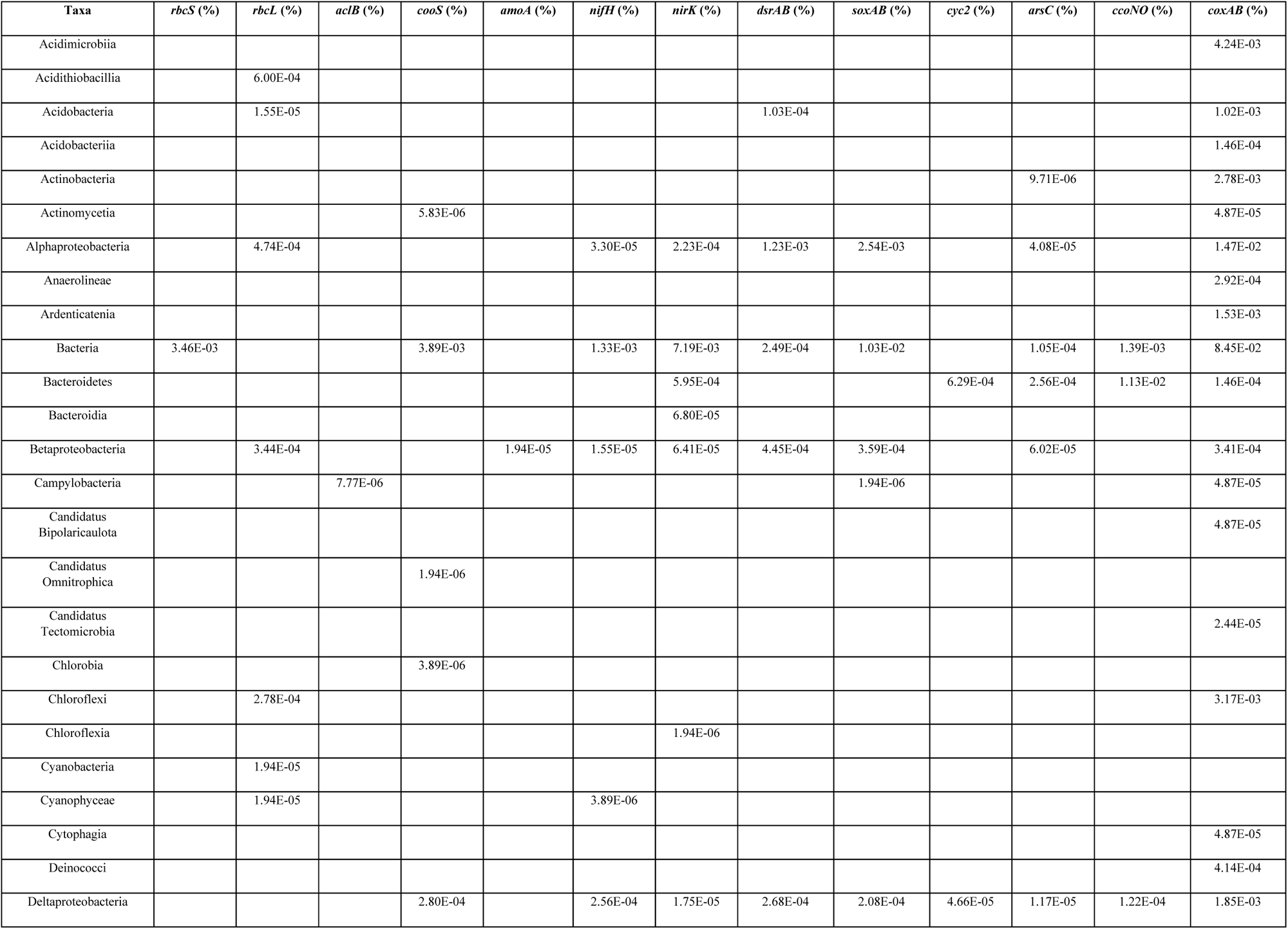
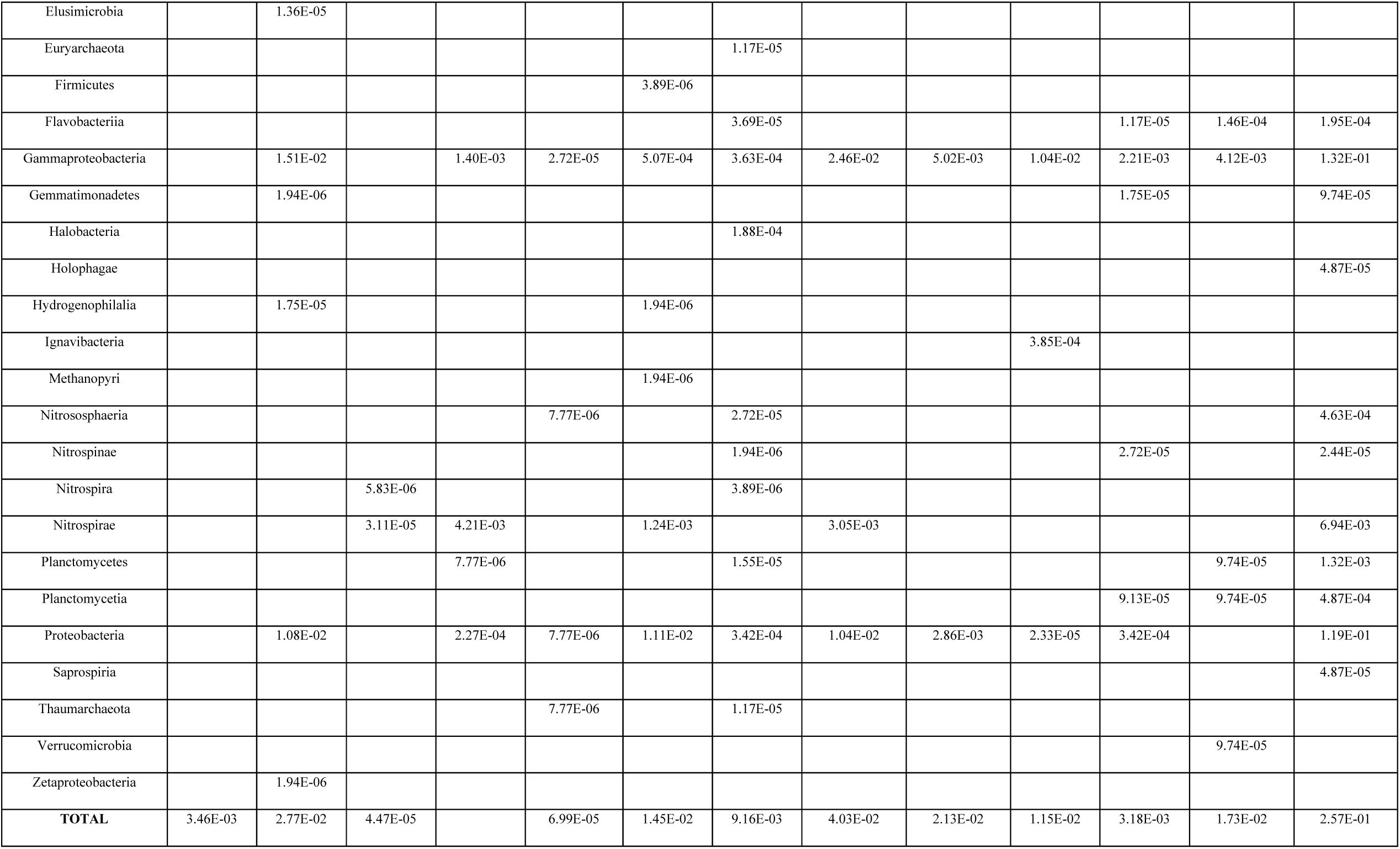
Relative abundance of different autotrophy genes and their taxonomic assignment for Castle Chimney.

**Supplemental Table 5.**
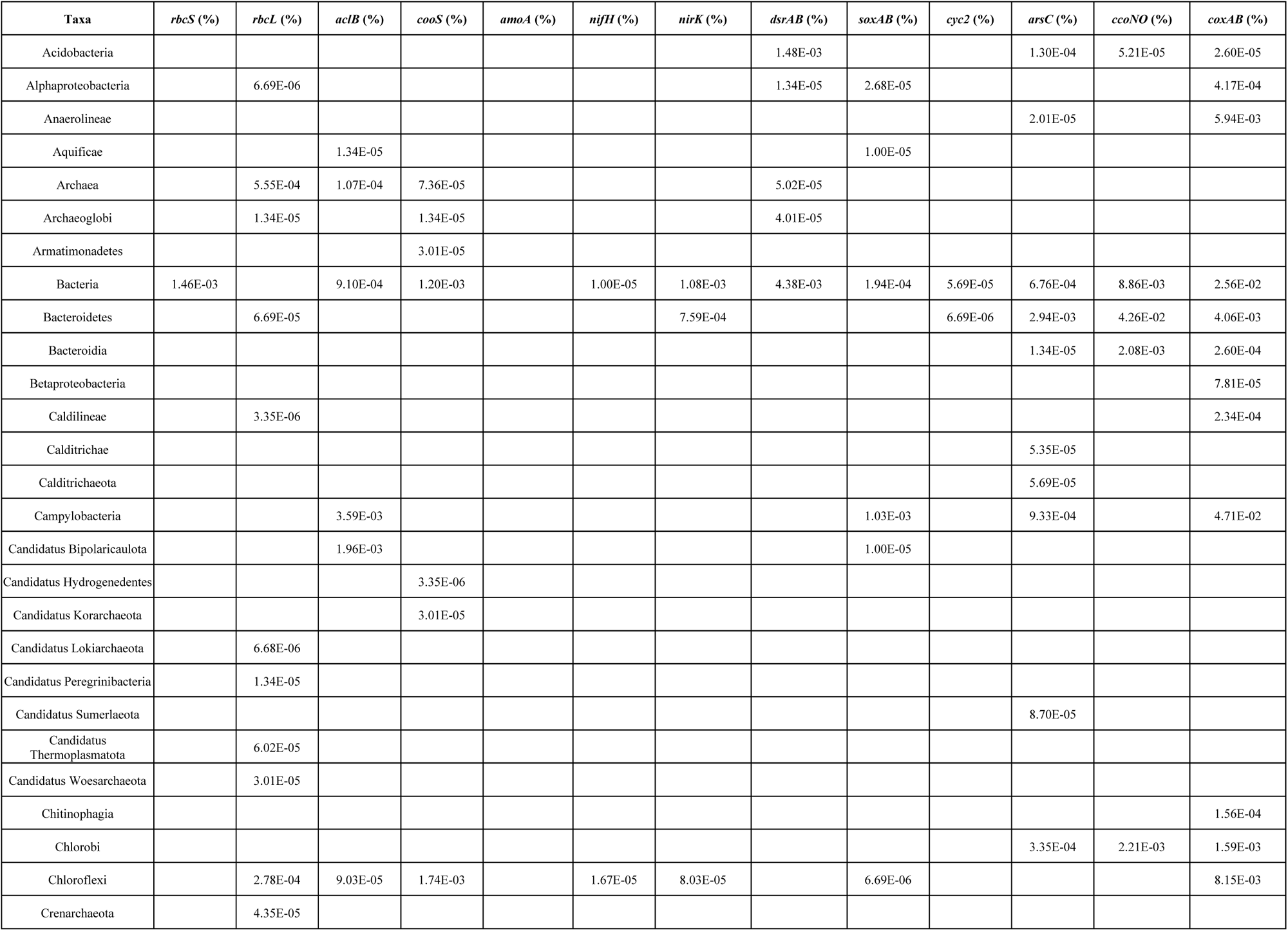
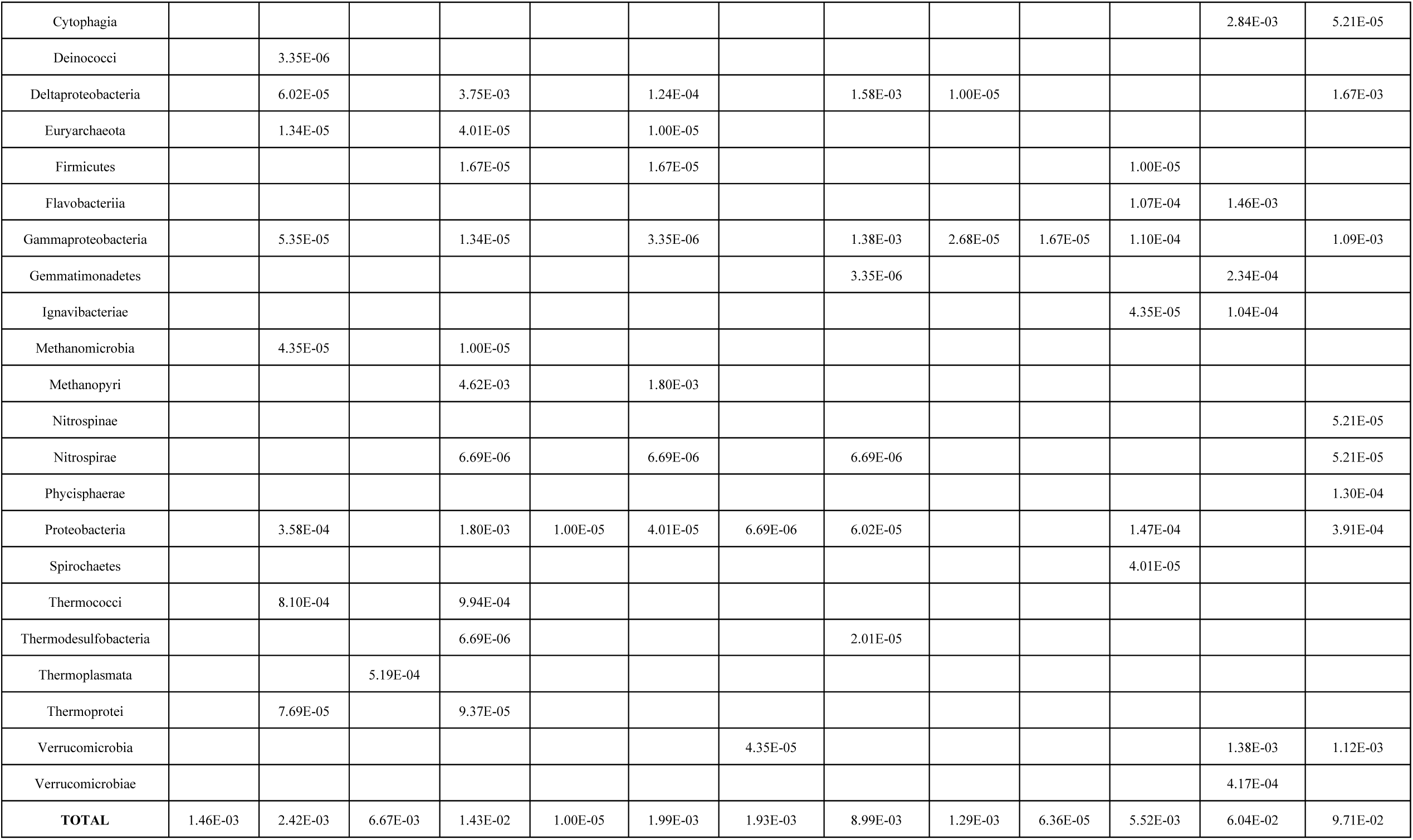
Relative abundance of different autotrophy genes and their taxonomic assignment for Pagoda Chimney.

**Supplemental Table 6.**
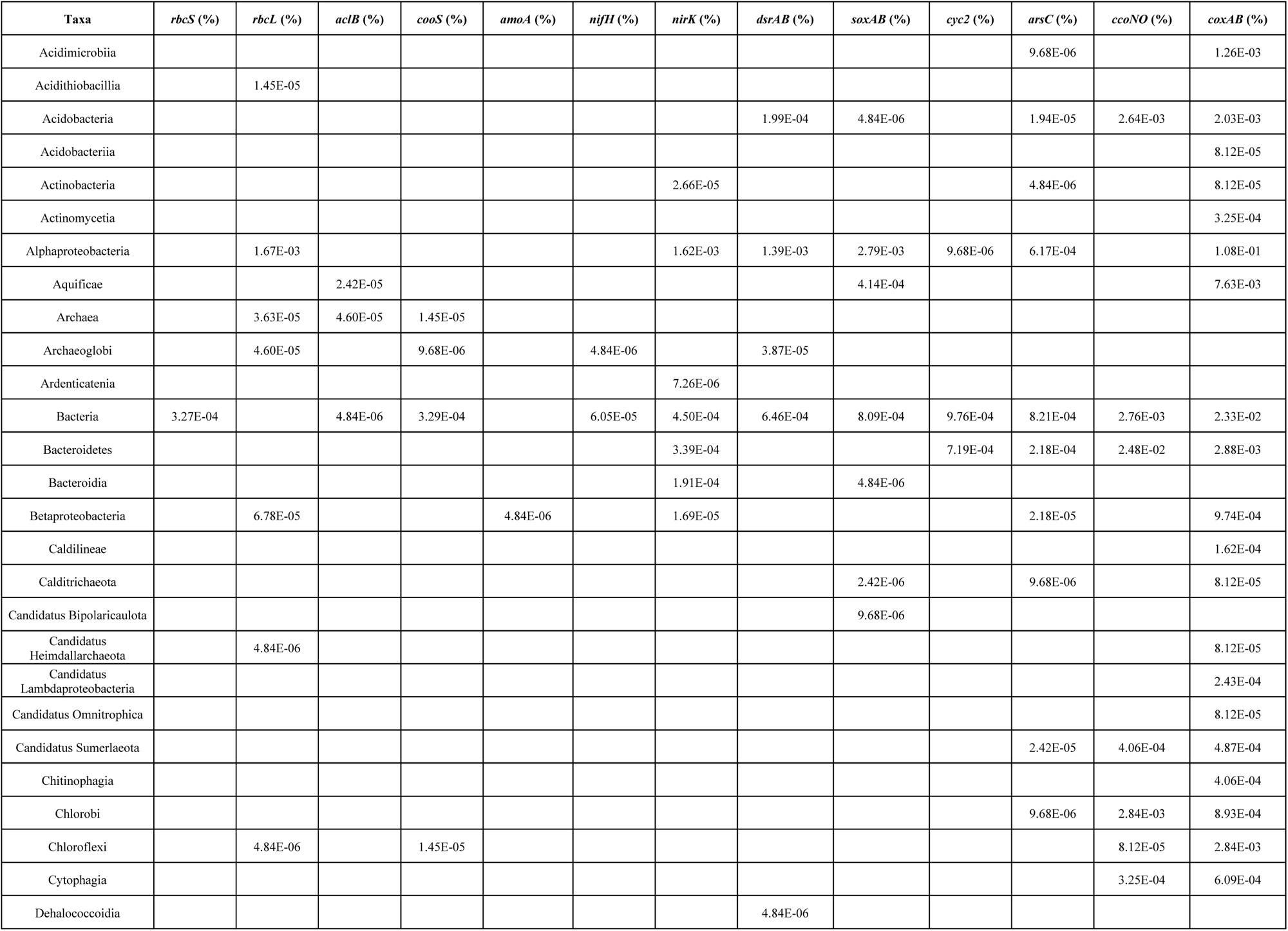
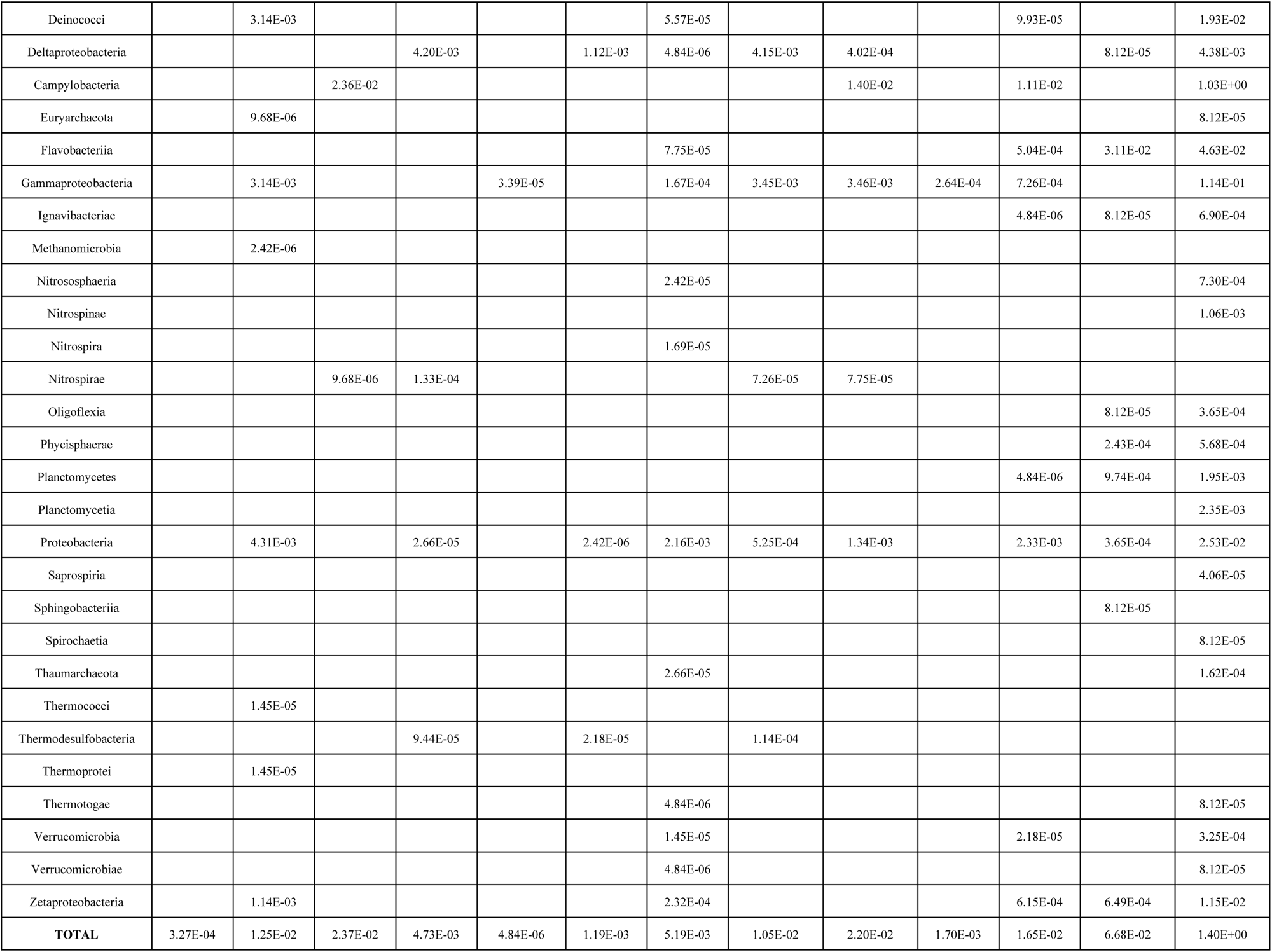
Relative abundance of different autotrophy genes and their taxonomic assignment for Snap-Snap Chimney.

**Supplemental Table 7.**
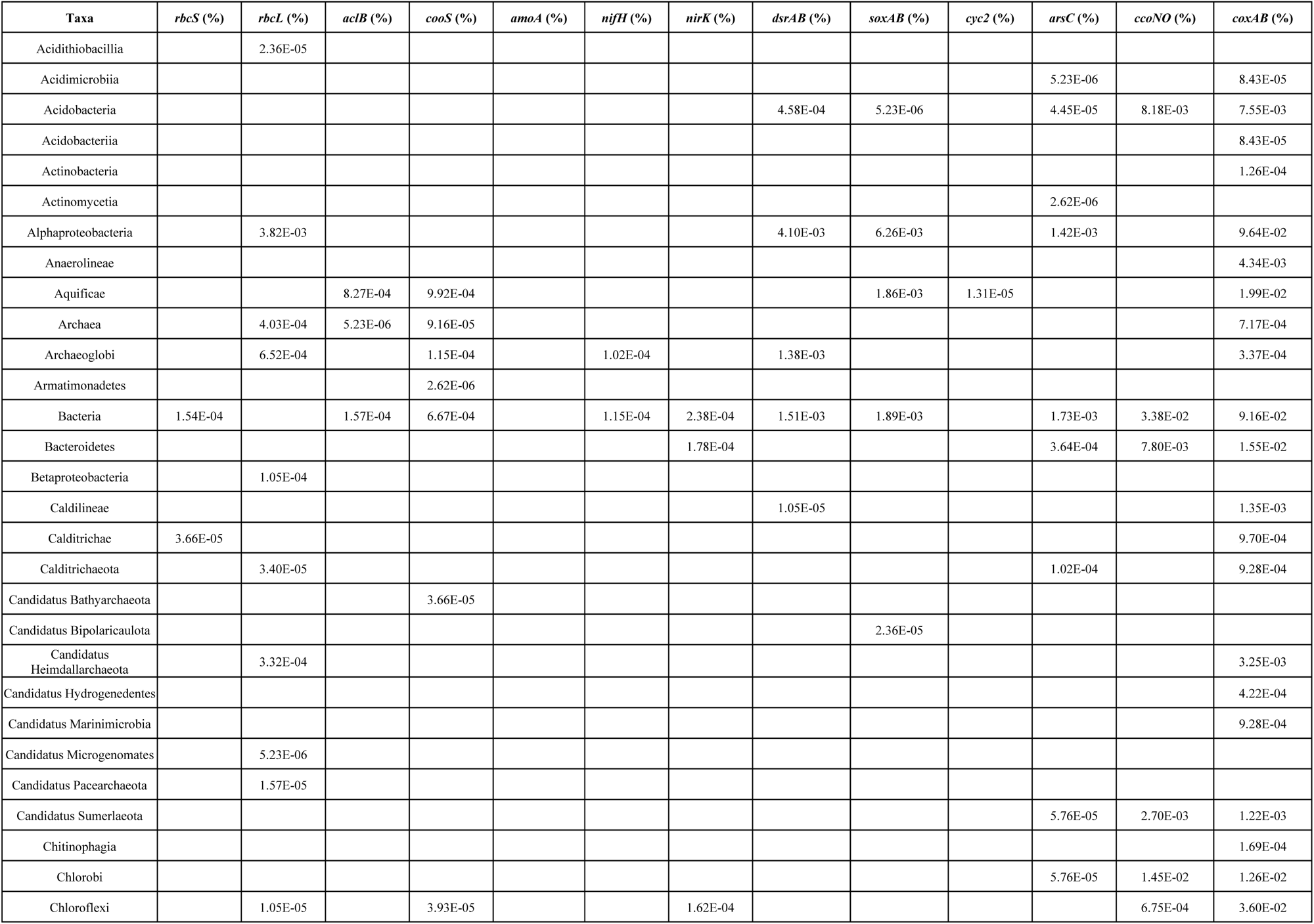
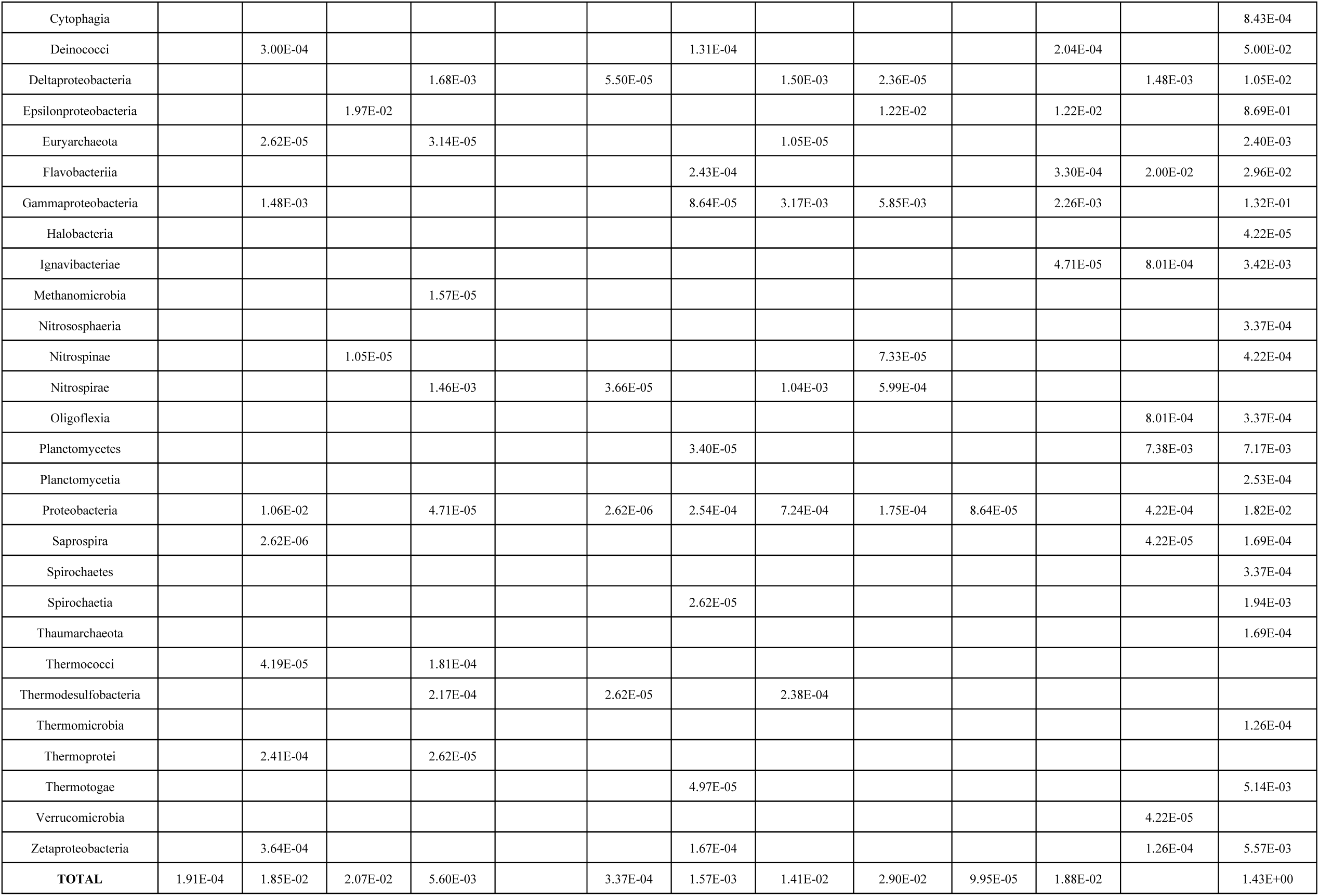
Relative abundance of different autotrophy genes and their taxonomic assignment for Ultra-No-Chi-Chi Chimney.

**Supplemental Figure 1.**
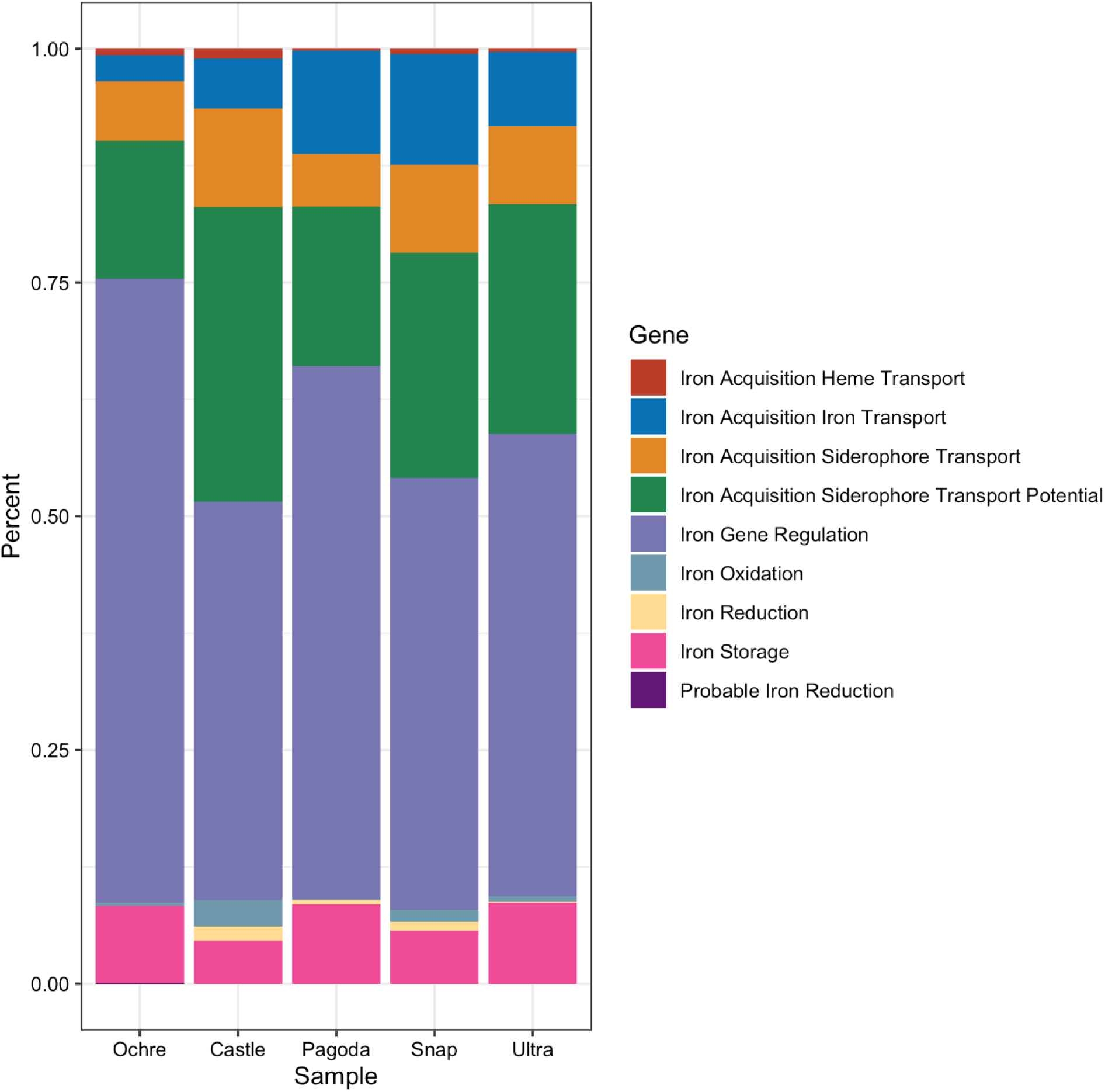
Stacked bar graph as a percentage of the whole of different types of iron genes found in each chimney from the FeGenie analysis.

